# Complete sequence verification of plasmid DNA using the Oxford Nanopore Technologies’ MinION device

**DOI:** 10.1101/2022.06.21.497051

**Authors:** Scott D. Brown, Lisa Dreolini, Jessica F. Wilson, Miruna Balasundaram, Robert A. Holt

**Author notes:** Corresponding Author - Robert A. Holt, 675 W 10th Ave, Vancouver, BC, Canada V5Z 1L3, Correspondence.

## Abstract

**Background:** Sequence verification is essential for plasmids used as critical reagents or therapeutic products. Typically, high-quality plasmid sequence is achieved through capillary-based Sanger sequencing, requiring customized sets of primers for each plasmid. This process can become expensive, particularly for applications where the validated sequence needs to be produced within a regulated and quality-controlled environment for downstream clinical research applications.

**Results:** Here, we describe a cost-effective and accurate plasmid sequencing and consensus generation procedure using the Oxford Nanopore Technologies’ MinION device as an alternative to capillary-based plasmid sequencing options. This procedure can verify the identity of a pure population of plasmid, either confirming it matches the known and expected sequence, or identifying mutations present in the plasmid if any exist. We use a full MinION flow cell per plasmid, maximizing available data and allowing for stringent quality filters. Pseudopairing reads for consensus base calling reduces read error rates from 5.3 % to 0.53 %, and our pileup consensus approach provides per-base counts and confidence scores, allowing for interpretation of the certainty of the resulting consensus sequences. For pure plasmid samples, we demonstrate 100 % accuracy in the resulting consensus sequence, and the sensitivity to detect small mutations such as insertions, deletions, and single nucleotide variants. In test cases where the sequenced pool of plasmids contains subclonal templates, detection sensitivity is similar to that of traditional capillary sequencing.

**Conclusions:** Our pipeline can provide significant cost savings compared to outsourcing clinical-grade sequencing of plasmids, making generation of high-quality plasmid sequence for clinical sequence verification more accessible. While other long-read-based methods offer higher-throughput and less cost, our pipeline produces complete and accurate sequence verification for cases where absolute sequence accuracy is required.

## Background

Sequence verification is essential for plasmids destined for downstream clinical research applications. Plasmids used for gene delivery (i.e. for generating genetically modified Immune Effector Cells), mRNA synthesis, or those used directly as plasmid DNA vaccines, must have their sequence identity confirmed prior to use as critical reagents or therapeutic products [1, 2]. Commercial services for clinical grade sequencing are available, but may offer longer than desired turnaround times and higher cost than some budgets may allow. A simple, fast, and inexpensive in-house option for plasmid sequence verification would be of potential utility. Here, we evaluate the Oxford Nanopore Technologies’ (ONT) USB-powered MinION sequencing device for this purpose.

Conventionally, capillary-based Sanger sequencing approaches have been used for plasmid sequence verification [3]. While these approaches can be inexpensive to run (around $5-10 per 800 bp read), the cost associated with outsourcing Sanger sequencing to a company offering a quality controlled environment with validated standard operating procedures is orders of magnitudes higher. Similarly, while Sanger sequencing is not technologically difficult to perform, it is onerous to set up and maintain in-house. For these reasons, another approach is desired. Long read sequencing is an attractive alternative as the sequence of the entire plasmid can be contained in a single read. Unlike plasmid sequence verification using short-reads [4, 5], long-read data is amenable to verifying repetitive regions, checking for inversions, and identifying other large-scale structural errors. However, long-read data generally suffers from higher error rates than short-read data, making the detection of small errors, such as SNVs, more challenging [6–8]. Despite these challenges, approaches have been developed to use ONT long read data to successfully sequence plasmids in a number of research applications [9–13], however, these existing tools and workflows are not appropriate for clinical research applications as they do not provide information on the confidence of each base call, generally do not provide quality measures for the resulting consensus sequence, and can introduce biases due to assumptions made when resolving ambiguous base calls that are the result of systematic errors [14]. In general, a pipeline compatible with downstream clinical research applications would be well-defined, validated, documented and controlled, and have clearly specified acceptance criteria that ensures the resulting sequence information is of at least the same quality that would be expected from the current gold standard of clinical grade Sanger sequence verification. Existing long read assemblers output a consensus sequence without any per-base quality metrics, and have difficulty processing the large numbers of reads required to gain confidence in the resulting sequence [15–17]. Recently, methods have been described that use ONT long read data to validate synthetic plasmid constructs [18, 19], focusing on the ability to multiplex plasmids into one sequencing run to reduce cost and increase throughput. While these methods have utility and value, they do not meet the requirements for high-quality plasmid sequence verification, such as in a clinical research setting, where a complete and error-free consensus sequence with interpretable quality scores is needed, and where multiplexing offers a potential source of contamination and uncertainty.

Here, we describe a cost-effective (compared to quality-controlled whole plasmid Sanger sequencing) and accurate plasmid library preparation procedure and analysis pipeline that generates a high-quality consensus sequence of the plasmid from an ONT MinION sequencing run, allowing for complete sequence verification of a plasmid prior to downstream applications. Utilizing the full capacity of the MinION flowcell, we obtain millions of raw reads for each plasmid which are then quality filtered to the subset of most accurate reads. We prioritize interpretability of the resulting sequence, providing confidence measures for each base of the consensus. We demonstrate that the resulting quality-filtered and processed consensus sequence achieves 100.00 % accuracy for three exemplar plasmids, and that mutated and contaminating templates can be detected at a level comparable to that of Sanger sequencing. Importantly, our entire process can be established to run within an individual lab, giving the operator complete control over all steps of the sequencing process.

## Results

### Oxford Nanopore Technologies MinION library construction and sequencing of linearized plasmid

Our novel approach was developed and validated using three unique plasmids previously generated in-house: BCRxV.TF.1, BCRxV.GagPolRev.1 and BCRxV.VSVG.1 [20] (**Additional file 1**). The sequence identity of each plasmid had been previously confirmed by Sanger sequencing following the NIH Human Genome finishing standard (> Phred30, double read coverage of every base) [21]. Prior to creation of the MinION sequencing libraries, we ran a restriction enzyme fingerprinting assay to verify the purity and identity of the plasmids (Materials and Methods).

Sequencing libraries were constructed using the SQK-LSK110 Ligation Sequencing kit following the Genomic DNA by Ligation protocol provided by ONT. Plasmids were linearized prior to library construction with restriction enzymes that generate 5’ overhangs. This is essential to maintain full plasmid sequence during the library construction end-repair step. We loaded each plasmid library onto a separate R10.3 MinION flow cell, and ran each flow cell for the full 72 hour run time. Each run generated an average of 3.86 million reads.

### Raw sequence base calling and read filtering

We performed base calling of the raw .fast5 data produced by the MinION using the GPU version of Guppy v5.0.11 on a GPU cluster (Materials and Methods). We used the ‘--min_qscore’ argument to filter reads by their lowest observed base quality score. We set this value to 12, only retaining base called reads where all bases had a quality score of 12 or higher. For our three plasmids, an average of 40.8 % of reads passed this quality filter (range 27.4 – 50.3 %). While the majority of reads were the expected sequence length for each plasmid (**Additional file 2: Figure S1**), we noted that a small subset of reads were dramatically shorter or longer than expected. These reads were all derived from the plasmid template, and may have arisen from fragmented template, incomplete transit of template through the pore, or concatenation of template molecules during library construction. We surmised that these reads of abnormal length would not benefit the downstream analysis, and only used reads with length within 250 bases of the expected template length. We observed between 70 – 87 % of quality-filtered reads meeting the length filtering criteria for use in subsequent analysis.

### Improving base calling accuracy by pseudopairing reads

Upon aligning these reads to the known reference sequence, we identified some cases where an incorrect base was consistently called predominantly in the forward or predominantly in the reverse read sets (**Figure 1**). This strand-specific bias may be the result of the specific sequence context within the pore at that position of the template, and differences in the ease that the base caller has in decoding the forward vs. reverse sequence motif [14, 22, 23]. Assessing the forward and reverse read sets independently, there was no way to determine which of the read sets contained the correct base call. When combining the forward and reverse read sets, there was a roughly equal number of reads supporting each base, resulting in the appearance of a heterozygous base call at that position. Therefore, another approach was needed to resolve this strand-specific bias.

**Figure 1.**
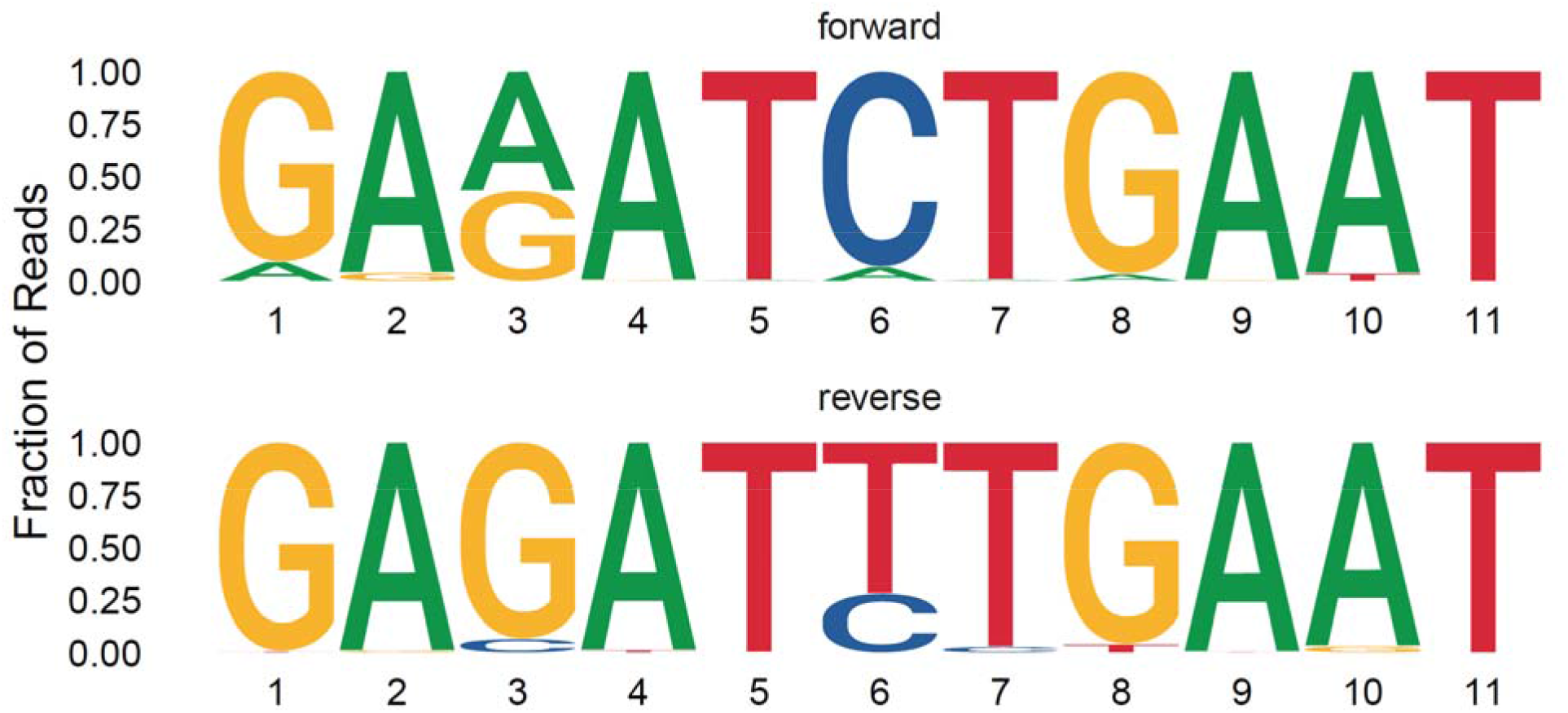
Strand-specific errors in base calling. Sequence logos are shown for one stretch of sequence from forward (top) and reverse (bottom) reads. The height of each letter gives the fraction of reads at that position that supported that base. Position 6 (and position 3, to a lesser extent) shows a strand-specific error – the correct base is C, but in the reverse reads a T is the most-evidenced base. When forward and reverse reads are pooled, there appears to be two bases at this position.

Paired base calling of matched forward and reverse strands of the same template molecule has shown great improvements in base calling accuracy [24]. This approach aligns the forward and reverse raw signals, and uses a consensus decoding algorithm to generate the base calls. This requires knowledge of which reads originated from the same template molecule. Generating such data requires special sequencing chemistry, such as ONT’s 2D or 1D^2^ kits, which were unavailable to us. Since we are sequencing a single plasmid, we know all reads should contain the same sequence. We hypothesized that we would be able to pseudopair forward and reverse reads to allow paired base calling of the raw .fast5 data, generating a single consensus duplex base call for every pseudopair of reads. We predicted that the consensus of the raw signal in these problematic strand-specific regions of the template would yield the correct base call. To generate these pseudopairs, we first aligned the Guppy base called reads to the expected reference as a way of annotating each read as coming from the sense or antisense strand of the template. Then, we paired sense and antisense reads, giving preference to pairs of reads with similar read lengths. These read pair lists were then used in conjunction with the raw .fast5 data as input into the Bonito base caller (https://github.com/nanoporetech/bonito; v0.4.0) to perform duplex base calling.

Overall, the Bonito duplex base called reads alleviated the strand-specific biases observed in the Guppy base called data. We noted, however, rare cases where miscalling was still occurring. Assuming that these miscalls were the result of modified bases [25], we used Bonito’s model finetuning feature to fine-tune the R10.3 base calling model using a random sample of 100 reads from one of our plasmids. This fine-tuning resulted in the elimination of systematic errors in our sequence reads for all three plasmids. Overall, our duplex base called reads had an order of magnitude reduction in error rate compared to the non-paired data (0.53 % compared to 5.3 % from the Guppy base called data, as reported by Qualimap).

### Generating a high quality plasmid consensus sequence

To obtain a consensus sequence for the plasmid from the high accuracy duplex base called reads, we implemented a “pileup” approach, taking the most-evidenced base at each position. To facilitate this, we generated a draft consensus by performing *de novo* assembly using a subset of the base called reads. We then aligned all reads to this assembly, producing an alignment matrix of all reads positioned in the same reference frame. This two-step procedure was more tractable, computationally, than performing a multiple sequence alignment of hundreds of thousands of reads. We then stepped through each position of the alignment matrix and enumerated the number of reads supporting each base. We handled insertions by tracking bases that aligned between two adjacent bases of the assembly. We visualized this data to get an overview of the strength of the signal and noise in our data (**Figure 2**).

**Figure 2.**
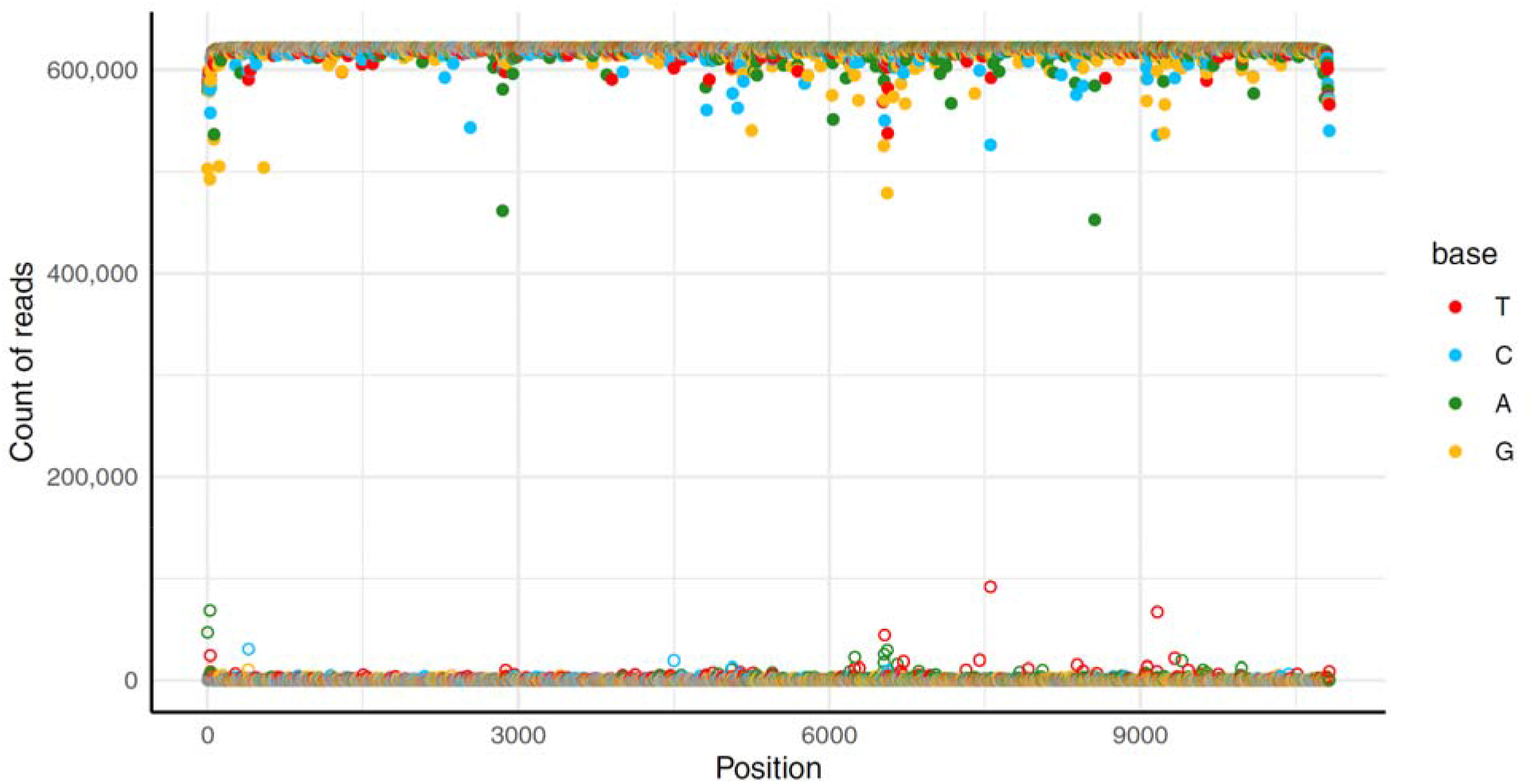
Overview of consensus read support across BCRxV.GagPolRev. Filled points denote the most-evidenced base at each position (x-axis) and hollow points denote the second-most-evidenced base. The y-axis shows the number of reads supporting each base. In general, nearly all reads support the top base, and there is good separation between signal (filled points) and noise (hollow points). For BCRxV.GagPolRev, the lowest signal to noise ratio is between the C and T at position 7554. We see a decrease in coverage at the extreme 5’ and 3’ ends of the plasmid.

Overall, we saw that the median signal:noise ratio across our three plasmids ranged from 3250 – 3770, while there were rare individual cases of it dropping to as low as 5.7. Generally, signal:noise ratios decreased at the extreme 5’ and 3’ ends of the template. Additionally, we observed drops in signal across homopolymer stretches (**Figure 3**). This is consistent with known difficulties in nanopore sequencing of homopolymer stretches [7], with reads underreporting the number of nucleotides in the homopolymer stretch. In one particularly long homopolymer from BCRxV.VSVG, only 17.3 % of the reads supported the full 17 bases. Using our data, an existing tool [18] was only able to report 15 of these 17 bases (data not shown).

**Figure 3.**
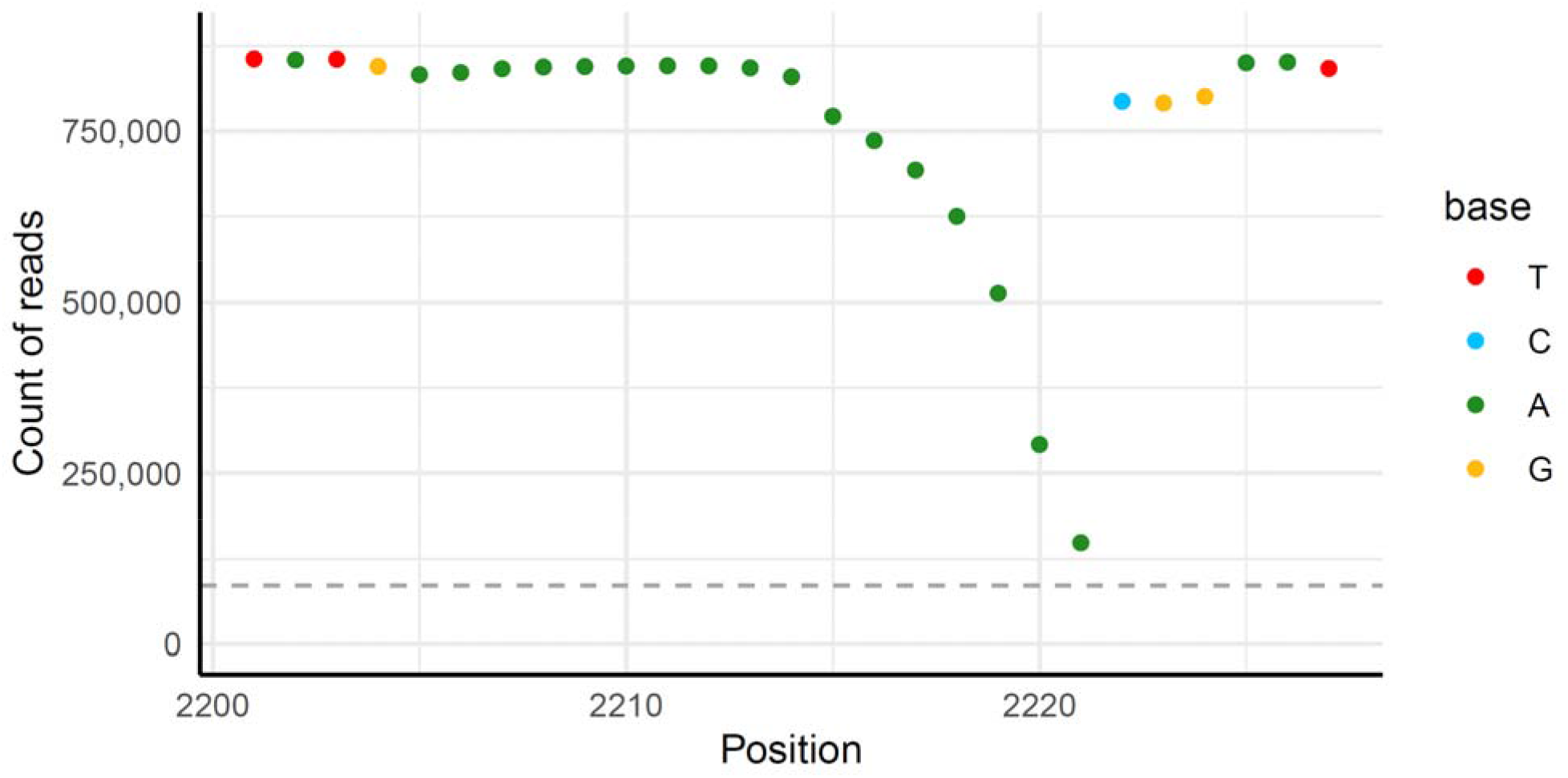
Decreased read support across homopolymer stretches. Points denote the most-evidenced base at each position across this subset of BCRxV.VSVG with a 17mer homopolymer. Due to difficulty base calling the exact number of bases in homopolymer regions, many reads report fewer than 17 bases. This results in an apparent coverage drop off across the homopolymer. The horizontal dashed grey line at 10 % of the max coverage represents the minimum base coverage required to add base to the consensus. Note that for this visualization, indels were right-aligned to show coverage dropping from left to right.

To generate a high quality consensus sequence from this data, we specified criteria that each base, and the overall consensus, must pass. We took the top-evidenced base at each position as the consensus base. The number of reads supporting this base must be at least 5× as high as the next most observed base at that position, and must be at least 10 % of the total number of aligned reads. Finally, the per-base confidence was calculated as the percent of reads at each position that support the consensus base, and the median per-base confidence across the entire plasmid must be greater than 99.9 %.

We observed rare cases where alignment artefacts near error-prone homopolymer regions resulted in two different consensus sequences depending on whether the *de novo* assembly reflected the sense or antisense strand (**Additional file 2: Figure S2**). In order to be confident that our resulting consensus sequence was accurate, we performed two alignment and consensus generation passes – once as described with alignment to the assembly, and once with alignment to the reverse compliment of the assembly (Materials and Methods). This yielded two consensus sequences, and if these sequences differ, this suggests that a variant template may be present (see next section, ***Detecting variant plasmids***).

### Detecting variant plasmids

Once we have obtained the high-quality consensus sequences (seeded from the sense and antisense versions of the assembly), we can determine if these sequences match the expected plasmid sequence. For our main goal of sequence verification, we simply compared the forward and reverse strand consensus sequences with the expected sequence (the “known” sequence of the plasmid based on cloning or synthesis designs) by sequence alignment and checked for any discrepancies. If any discrepancies were found in either consensus, we reported that the sequenced plasmid contains that variant. In the case where any position of the consensus did not meet our criteria, an “N” base is added to the consensus. The base signal strengths and confidence scores for these positions should be manually inspected, as they may be the result of a decreased signal to noise ratio in a difficult to sequence area, or could be the result of a variant in the plasmid.

As a secondary goal, we can also detect the presence of subclonal templates in the population of plasmid molecules used in library construction. It is rare that such a case would arise, but could exist if a mutation occurred early within the bacterial culture’s growth phase, or the bacterial culture was seeded from two bacteria in close proximity. Based on the thresholds we have set for consensus sequence generation, we expected a theoretical lower limit of detection for subclonal variant templates to be 20 % for single nucleotide variants (SNVs) and insertions (a variant template present at a frequency of less than 20 % would be indistinguishable from noise). For deletions, since we allowed coverage of a single base to drop to as low as 10 % of all aligned reads (to detect all bases within homopolymers), we expected that the sensitivity to detect subclonal deletions would be low, requiring greater than 90 % of the templates to contain the deletion. We tested these expectations, experimentally, by designing a mutant version of BCRxV.VSVG containing three SNVs, three insertions, and three deletions (**Additional file 1**). This variant plasmid was sequenced as described for the other plasmids, and the raw .fast5 reads were mixed with raw wildtype BCRxV.VSVG reads *in silico* at varying frequencies, allowing random pseudopairing of wildtype and mutant reads to occur as they would have if the templates were mixed, physically, prior to sequencing. The analysis was run for each dataset, which demonstrated that the sensitivity for detecting subclonal variant templates was consistent with our predictions. Two of the three insertions were detected when their template frequency was approximately 20 %, with the other (which resulted in a lengthening of the 17mer homopolymer, and so would be predicted to be harder to detect) was detected at 40 % frequency (**Figure 4**). The three SNVs were detected with template frequencies ranging from 25 – 30%. The deletions were not detectable until the variant template was at a frequency of 95 %. At lower variant template frequencies, deletions were indistinguishable from the drop in signal observed within homopolymer regions, and are not able to be reliably detected.

**Figure 4.**
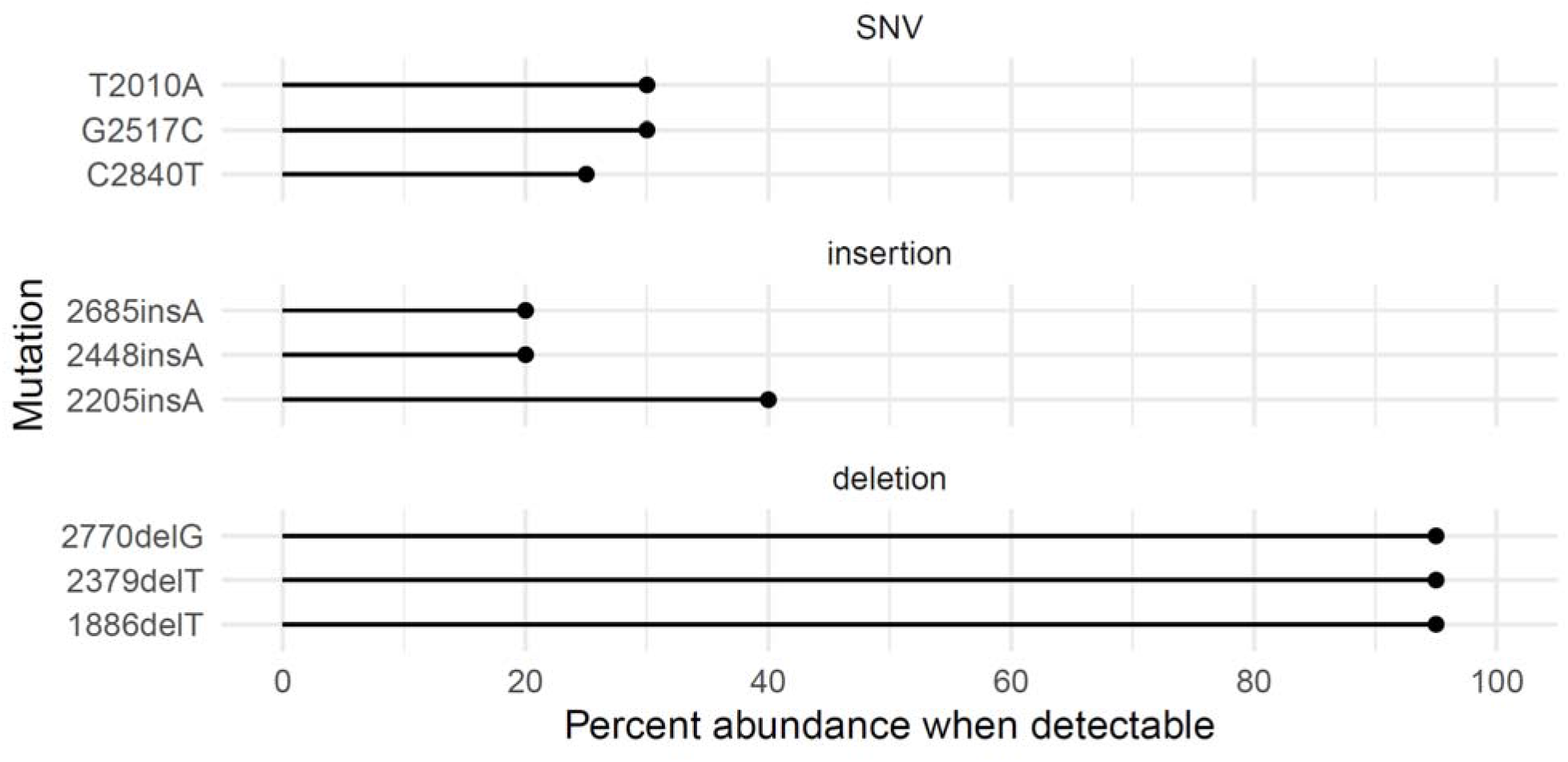
Subclonal variant template detection sensitivity. For three different mutation classes (SNV, insertion, deletion), the minimum percent abundance of variant reads (x-axis) that allowed detection of the mutation in the *in silico* generated mixed datasets is plotted. SNVs and insertions are detectable when around 30 % of reads are from the variant template, while deletions were not detectable until 95 % of the reads were from the variant template.

Contaminating, unrelated plasmids are another type of variant template that we may wish to detect in our sample. These are plasmids whose presence should be detected by fingerprinting assays performed prior to sequencing; however it is possible, though exceedingly unlikely, that an unrelated plasmid would have the same fingerprint and length as the expected plasmid. As reads from such a plasmid would not be expected to align well to the expected reference, we can count the fraction of reads which are successfully pseudopaired, since this step only uses reads which have a single continuous alignment to the expected reference that is within 250 bp of the reference length. In our three sequencing runs, 92.3 – 97.2 % of quality- and length-filtered reads were successfully pseudopaired. In a simulation where we ran our pipeline using a subset of 10,000 quality- and length-filtered reads, but used a length-matched incorrect reference, 0 % of the reads were successfully pseudopaired. Hence, this metric can be checked to ensure there is no significant contamination of an unrelated plasmid, with values less than 90 % being evidence of plasmid contamination.

### Cost analysis

We have described a sample preparation and analysis pipeline that has comparable performance to outsourcing Sanger sequencing to a commercial provider offering services acceptable by regulatory bodies. While we have comparable performance, our method results in significant cost savings to the user. At the present time, consumables cost for our method includes the ONT library preparation kit (~$160 CAD per sample), the ONT MinION flowcell (~$1,200 CAD per sample), and electricity to run the GPU-powered basecalling (eight RTX 3090 GPUs [at 450 Watts of power each] × five days = 432 kWh, as an upper bound assuming 100% usage), which at ~$0.10/kWh in British Columbia, Canada, totals ~$50 CAD. Adding other miscellaneous consumable costs associated with sample preparation, our method totals ~$ 1,500 CAD per plasmid. Note that there will be additional costs (maintenance, administration, labor), but as these costs can vary greatly by institution, we have not included them here. Based on quotes we have obtained from commercial Sanger sequencing providers, sequencing a ~10 kb plasmid to meet regulatory standards would cost ~$15,000 CAD per plasmid, or 10× the cost of our method. One reason the cost for clinical grade sequencing is so high is because primer sets used for primer walking are specific for each plasmid template and must be pre-validated. Even if a user needed to purchase the MinION device (~$1,200 CAD) and a GPU compute server (~$35,000 CAD), our method is potentially cheaper than outsourcing after only three sequence verification runs (~$40,700 CAD vs. ~$45,000 CAD when outsourcing).

## Discussion

We have described a pipeline that generates a high-quality consensus sequence of linearized plasmid using ONT MinION sequencing, leveraging substantial sequencing depth and stringent quality filters to overcome the relatively high error rates associated with nanopore sequencing. The purpose of this method is to verify the identity of a pure sample of plasmid with known sequence. For pure samples of linearized plasmid, our pipeline produces the true sequence with a median per-base confidence of over 99.9 %.

Our pipeline improves typical read error rates by an order of magnitude by performing paired base calling on pseudopaired reads, leveraging the fact that we are sequencing a single plasmid. True duplexing would only improve this result, whether from a technological advance to generate paired data natively at the time of sequencing, or barcoding of every template molecule with a unique molecular identifier (UMI) prior to sequencing. This would ensure that paired reads came from the same template molecule, and would increase the sensitivity for detection of subclonal variant templates.

This pipeline is first and foremost a sequence verification method. In this embodiment, the user must know the expected sequence of the plasmid. Due to this, the approach is not entirely unbiased. Additionally, we report a single consensus sequence of all plasmid templates present in the sample rather than separate consensus sequences for each plasmid, if there should be more than one. Future developments could extend this framework to be run completely independently of the expected reference sequence, allowing for multiple consensus sequences to be output if multiple templates are detected.

With our current pipeline, subclonal insertion and SNV variants are detectable when they are at roughly 25 % frequency. Due to the necessarily low coverage threshold needed for homopolyer stretches (at least 10 %), our sensitivity to detect subclonal deletions is low. It is conceivable that one could set a more stringent coverage threshold (80 %) globally while having a relaxed threshold (10 %) specifically within homopolymer regions. This would increase the sensitivity of non-homopolymer deletions to be comparable with insertions and SNVs, however, deletions within homopolymers would still require greater than 90 % frequency to be detected. For the current implementation, we acknowledge this limitation in detecting deletions.

Homopolymers themselves present challenges for accurate base calling. Initially, we tested R9.4.1 flowcells, but found the performance within homopolymer regions to be insufficient. R10 generation flowcells are known to improve accuracy within homopolymers, increasing the number of nucleotides that are read within the pore at once [26]. There is certainly an upper limit to the length of a hompolymer that can be accurately sequenced using our approach, though we have not yet identified it. Within our test plasmids, we have demonstrated successful sequence determination for homopolymers up to and including an 18mer.

To our knowledge, our pipeline is the first to use long read sequence data to sequence verify plasmids at a level consistent with quality controlled and regulated environments. We summarize our method in the context of other related methods in **Table 1**. Currin et al. [14] describe a highly multiplexed method with a unique approach to dealing with strand-specific sequencing errors, analyzing forward and reverse reads separately in a statistical framework to identify the correct base. However, their method is reference-based, and defaults to the reference base in regions with unreliable sequencing data. Recent work by Emiliani et al. [18] describe a tool called Circuit-Seq which is also multiplexed, and utilizes a standard barcode-splitting, assembly, and polishing workflow. Circuit-Seq only outputs a consensus sequence without any per-base quality metrics, and has an initial error correction step which has the potential to introduce biases or decrease sensitivity. A pre-print by Mumm et al. [19] describes OnRamp, which is also primarily an assembly and polishing procedure. OnRamp produces consensus sequences with a quality score based on the number of variants contained in the consensus, but no per-base quality measures. Our method is distinct from these methods in that we pseudo-pair reads prior to basecalling to yield higher quality base calls, and instead of polishing an assembly, we align reads to a draft assembly and use the read alignment to, position by position, infer the consensus base and enforce quality filters. Rather than simply outputting a .fasta file with the consensus sequence, we can interrogate any position to determine how confident the consensus base is, or if there is evidence of subclonal variants present, making this more comparable to traditional Sanger sequencing workflows. This is crucial for any downstream clinical research applications of these plasmids, as sequence verification is required by regulatory bodies.

**Table 1.**
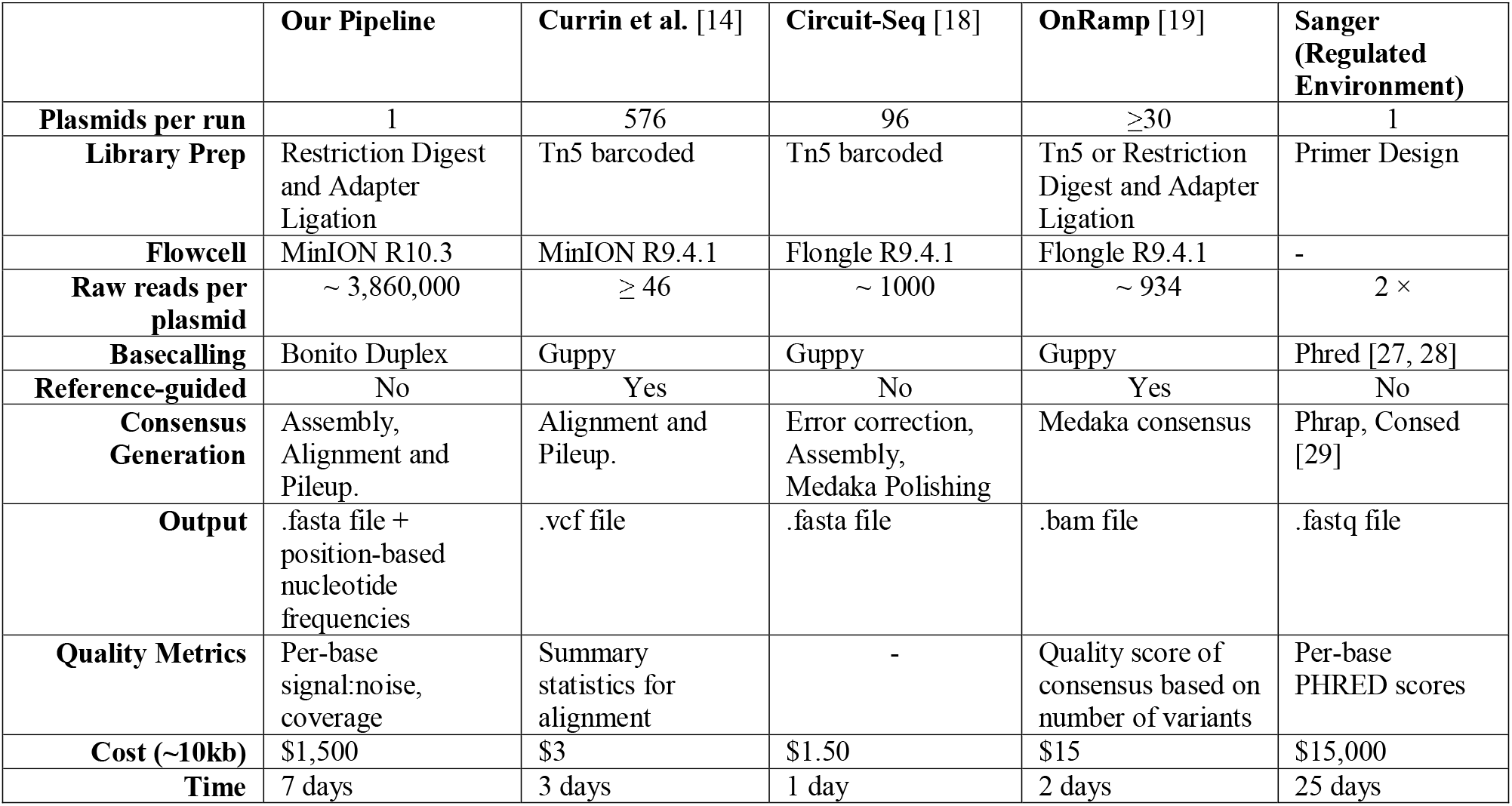
Comparison of plasmid sequence verification methods.

We have described a pipeline to sequence verify plasmids at a level appropriate for clinical research applications with a 90 % reduction in cost. This pipeline uses a full flowcell per plasmid, eliminating any possibility of cross contamination between plasmids either barcoded and pooled, or plasmids run sequentially on the same flowcell. If this pipeline were to be used in a more relaxed setting, MinION flowcells could be reused by running a sample for less time, washing, and then reloading the flowcell with a new sample. This would allow multiple plasmids to be run on the same flowcell, reducing overall cost. Alternatively, barcoding independent plasmids prior to sample loading would allow multiple plasmids to be run at the same time, also reducing costs, as has been done for the related methods described above [14, 18, 19]. While we have benchmarked this procedure on R10.3 flowcells, we fully expect that future developments to this platform (R10.4 and beyond) will only improve the accuracy. Before running this analysis on R10.4+ data, it is likely that the base calling model fine-tuning would need to be repeated for the appropriate flowcell-matched base calling model. Alternatively, since the fine-tuning of the base calling model was likely required due to base modifications such as methylation, ensuring the plasmid is free from modified bases prior to library construction may obviate the need to fine-tune the model. It is also important to note that the feasibility of our plasmid sequencing approach is ultimately reliant on the availability of compatible flowcells from ONT. At the time of writing, ONT is rapidly iterating on their R10 generation flowcells, and thus availability is unpredictable. With each change to flowcell or related chemistry, this sequencing and analysis procedure must be re-validated to ensure the performance is not decreased from previous versions. It is our understanding that the R10 generation will become the stable product line once it is finalized. We have made all code in the pipeline available (**Additional File 3** and https://github.com/scottdbrown/minion-plasmid-consensus) so that users may adapt and update the pipeline and models to use the currently available flowcells. This also allows users to make any modifications to the code to run on the specific compute architecture available to them.

## Conclusions

The major benefit of our pipeline over assembly and polishing approaches is that the output of our pipeline includes, in addition to a consensus sequence, the underlying per-base data for each position of the consensus. This is somewhat analogous to traditional Sanger sequencing approaches used for clinical sequence verification, and allows for manual inspection and interpretation of the quality of the resulting consensus. This allows the confidence, per base, to be assessed, strengthening the ability to use this pipeline as a replacement for Sanger sequencing of plasmids for clinical research use.

Our pipeline can provide significant cost savings compared to outsourcing clinical-grade Sanger sequencing of plasmids, with estimated costs an order of magnitude lower for a single plasmid. For labs with access to a MinION sequencer and compute infrastructure, this approach could make generation of high-quality plasmid sequence for clinical sequence verification more accessible.

## Materials and Methods

### Restriction enzyme fingerprinting assay

The purity of each plasmid sample to be sequenced was confirmed by restriction enzyme fingerprinting prior to library construction. A pair of restriction enzymes was chosen to generate a unique banding pattern for each construct. Reactions were set up with 2.5 μg plasmid, 180 total units of restriction enzyme (New England Biolabs), plus 1 unit of Topoisomerase I (New England Biolabs, M0301S) to promote complete digestion. Digestions were run according to the manufacturer’s instructions. The resulting banding patterns were visualized on 0.8% E-gels (Thermofisher, G501808). A representative gel is shown in **Additional file 2: Figure S3**.

### Sequence library construction

Plasmids were grown in NEB® Stable Competent *E. coli* (New England Biolabs), which are *dam^+^/dcm^+^.* Plasmids were prepped using PureLink™ Expi Endotoxin-Free Giga Plasmid Purification Kit (Thermofisher, A31233). Six-hundred fmol of each plasmid was linearized with a restriction enzyme (New England Biolabs; BCRxV.TF: AscI, BCRxV.VSVG: BamHI, BCRxV.GagPolRev: BamHI) chosen to generate a single cut site with 5’ overhangs. Each reaction was supplemented with 1 unit of Topoisomerase I (New England Biolabs, M0301S). Complete digestion was confirmed by running 200 ng of undigested plasmid alongside 200 ng of digested sample on 1.2% E-gel (Thermofisher, G21801). Samples were purified using Qiaquick PCR Cleanup kit (Qiagen, 19086) following the manufacturer’s instructions, and eluting with 50 μL pre-warmed (50°C) EB (Qiagen, 19086).

Libraries were prepared using the ONT SQK-LSK110 kit following the Genomic DNA by Ligation protocol (GDE_9108_v110_revC_10Nov2020). Samples were quantified with Qubit dsDNA BR kit (Q32850). 300 fmol of linearized plasmid was repaired using NEBNext FFPE DNA Repair Mix (NEB, M6630S) and NEBNext Ultra II End Repair/dA Tailing Module (NEB, E7546S). Samples were purified with a 1× volume of Ampure XP beads (Beckman, A63880), and eluted with 61 μL pre-warmed (50°C) Qiagen EB. Adapter ligation was performed with NEB Quick Ligase (E6056S), ONT Ligation Buffer (LNB) and ONT adapter mix (AMX-F). Samples were purified with 0.5× Ampure XP beads, and washed with ONT Long Fragment Buffer, and eluted with 15 μL ONT Elution Buffer (included in SQK-LSK110 kit).

### Sequencing using ONT MinION

MinION R10.3 (FLO-MIN111) flowcells were prepared and run according to ONT instructions: 50 fmol of prepared library was loaded per flowcell, and the run time was set to 72 hours. The MinION USB device was connected to a computer running Windows 10, with an Intel Core i7-9700 8 core CPU, 32GB RAM, and a 1TB SSD data drive. On-device base calling was disabled, instead performing base calling in a later step on our higher-powered computational resources.

### Base calling raw sequence data with Guppy

We assembled all steps of our pipeline into a Snakemake v5.7.4 workflow (**Additional file 2: Figure S4**). We performed initial base calling of the raw .fast5 data using the GPU version of Guppy v5.0.11. We ran base calling on a GPU cluster comprised of eight NVIDIA GeForce RTX 3090 GPUs, 256GB RAM, and two Intel Xeon Silver 4212 12 core CPUs. Each .fast5 file generated by MinKNOW was submitted to the cluster in its own job, requesting 12GB RAM and 1 GPU. We used the “dna_r10.3_450bps_hac.cfg” config file and “template_r10.3_450bps_hac.jsn” model file packaged with Guppy, and set “--min-qscore” to 12. Base calling took roughly 1 minute per .fast5 file. We used a custom Python script to filter the resulting base called reads to those that had length within 250bp of the expected sequence length.

### Pseudopairing reads

In order to pseudopair forward and reverse reads, we performed an initial alignment of the base called reads to the expected reference sequence. We used minimap2 v2.17, with the “-x map-ont” and “--cs” options, generating a .paf file. We then use a custom Python script to parse this .paf file, generating a list of forward and reverse read names which contain only a single alignment to the reference, and for which the number of aligned bases is no shorter than the reference length minus 500 bases. We sort these two lists by read length, and step through each simultaneously, outputting pseudopaired forward and reverse read names to a file.

### Fine-tuning the Bonito base calling model

We performed Bonito base calling model fine-tuning as described in the Bonito manual. We fine-tuned the “dna_r10.3” model included with Bonito v0.4.0 using 100 reads of our BCRxV.VSVG plasmid, requesting 64GB of RAM and using a batch size of 32, 1 epoch, and a learning rate of 5e-4. We used this fine-tuned model for all Bonito duplex runs. We have made the fine-tuned model available at https://doi.org/10.5281/zenodo.6626041.

### Paired base calling with Bonito

We used a modified version of Bonito v0.4.0 to perform the duplex base calling of the read pairs, making some minor adjustments to the Bonito code to prevent the process from using too much memory on our system (https://github.com/scottdbrown/bonito/releases/tag/v0.4.0a). We submitted the Bonito duplex base calling jobs to our GPU cluster, requesting 40GB of RAM and 6 CPUs per job. We set “--max-cpus” to 4, and used our own fine-tuned base calling model. On average, Bonito duplex base calling jobs of around 750 read pairs took 20 minutes to complete.

### De novo assembly to generate a reference

We combined all duplex base called reads into a single .fasta file. We subsampled 500 reads from this file to use for *de novo* assembly using Canu v2.0 [16]. We set the “-corrected” and “-nanopore” options, set the “genomeSize” to the length of our expected reference, and used 16 threads. We extracted the first sequence in the resulting “unitigs.fasta” file to use as the reference sequence for subsequent read alignment and consensus sequence determination.

### Consensus sequence generation

To generate a consensus sequence for our sequence data, we align the duplex base called data to the *de novo* assembled reference sequence. Because of the logic of our consensus sequence building, and due to the standard practice of left-aligning gaps in indels, we perform two separate consensus sequence determination steps using both the *de novo* assembly as provided, as well as the reverse compliment. This is to ensure that any variant near a homopolymer stretch does not have the variant support diluted due to alignment artefacts (**Additional file 2: Figure S2**).

We align the duplex base called reads to the *de novo* assemblies, separately, using minimap2 v2.17, with the “-x map-ont” and “--cs” options, generating a .paf file. We then use a custom Python script to parse the alignments. We build an array with positions for 5’ of the reference sequence, every base of the reference sequence, every position in between bases of the reference sequence, and 3’ of the reference sequence. We step through every read alignment and update a dictionary at each position recording the number of reads supporting each observed base at each position (with multiple bases represented as an array of dictionaries). We then build the consensus sequence by stepping through every position of the array and adding the most evidenced base if it meets our criteria: (1) the top base being at least 5× more abundant than the second top base, and (2) the coverage of the top base being at least 10 % of the max coverage. If only the first criterion is not met, an N base is added to the consensus, signifying insufficient read support to accurately call a base at this position. If the second criterion is not met, no base is added to the consensus.

### Variant plasmid design

To test the detection of subclonal variant templates, we designed a 1040 bp variant insert for our BCRxV.VSVG plasmid with the following nine mutations: 1886delT, T2010A, 2205insA, 2379delT, 2448insA, G2517C, 2685insA, 2770delG, and C2840T (positions given relative to resulting sequence after linearization with BamHI) . We had this fragment synthesized by Azenta Life Sciences, and cloned it into our wildtype BCRxV.VSVG plasmid using SfuI and BalI restriction sites.

### Subclonal variant simulation

We sequenced our variant BCRxV.VSVG plasmid as described for the other plasmids. To simulate sequence datasets with BCRxV.VSVG variant plasmids at varying levels of abundance, we first performed Guppy base calling on all wildtype and variant data to obtain the read names that passed quality filtering. We then created raw .fast5 read sets of 10,000 total reads *in silico*, ranging from 0 % to 100 % variant reads in 5 % increments. We then processed these read sets as described, allowing random pseudopairing of wildtype and variant reads to occur, and interrogated the resulting consensus sequences to determine at what variant read frequency each mutation was detected.

### Quality reports

To obtain quality metrics of our duplex base called reads, we used Qualimap v2.2.2 to provide a summary report on these reads aligned to the expected reference sequence. We used minimap2 v2.17 with the “-ax map-ont --cs” options to generate a .sam file, used samtools v1.7 to convert the .sam file to a .bam file and to index this .bam file, then ran Qualimap bamqc with 32 threads and 16GB of memory. From this report, we obtain the number of aligned reads, average coverage, mean error rates of the reads, and view the plots to ensure there are no issues with the data.

## List of Abbreviations

ONT: Oxford Nanopore Technologies
SNV: Single Nucleotide Variant

## Declarations

### Ethics approval and consent to participate

Not applicable.

### Consent for publication

Not applicable.

### Availability of data and materials

Custom code is available at the following GitHub repository: https://github.com/scottdbrown/minion-plasmid-consensus. All files found in the GitHub repository at the time of publication can also be found in Additional File 3. Raw .fast5 sequence data for our four sequencing runs is available at SRA under Bioproject PRJNA905101 (https://www.ncbi.nlm.nih.gov/bioproject/PRJNA905101).

### Authors’ Contributions

RAH and SDB designed research, LD performed plasmid preparation, sequence library construction, and plasmid sequencing, SDB analyzed data and developed the pipeline, JFW and MB oversaw quality control and data interpretation, and SDB, LD, and RAH wrote the paper. All authors read and approved the final manuscript.

## Acknowledgments

We would like to thank Luka Culibrk for helpful review of the code.

