## Additional File 2 for "Complete sequence verification of plasmid DNA using the Oxford Nanopore Technologies’ MinION device"

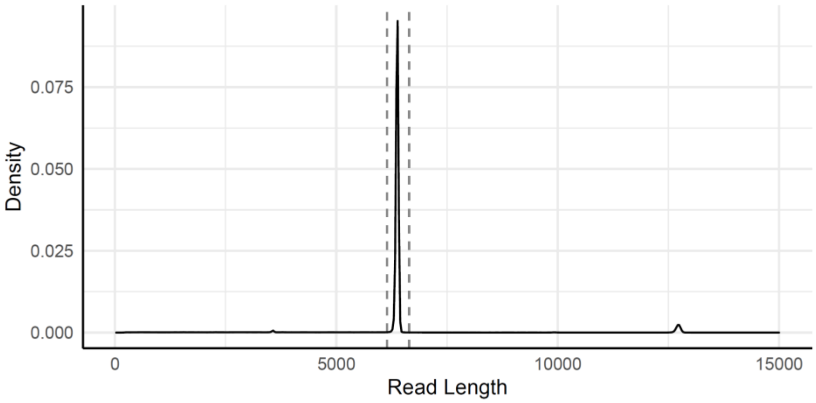


**Figure S1. Raw read length distribution for BCRxV.VSVG.** A density plot showing the distribution of quality-filtered read lengths obtained from BCRxV.VSVG. The region between the dashed vertical lines show the read lengths that pass length-filtering (are within 250 bp of the expected reference length). The x-axis has been truncated at 15,000 bp, removing 0.4 % of reads with length longer than 15,000 bp. The longest observed read had a length of 356,764 bp.

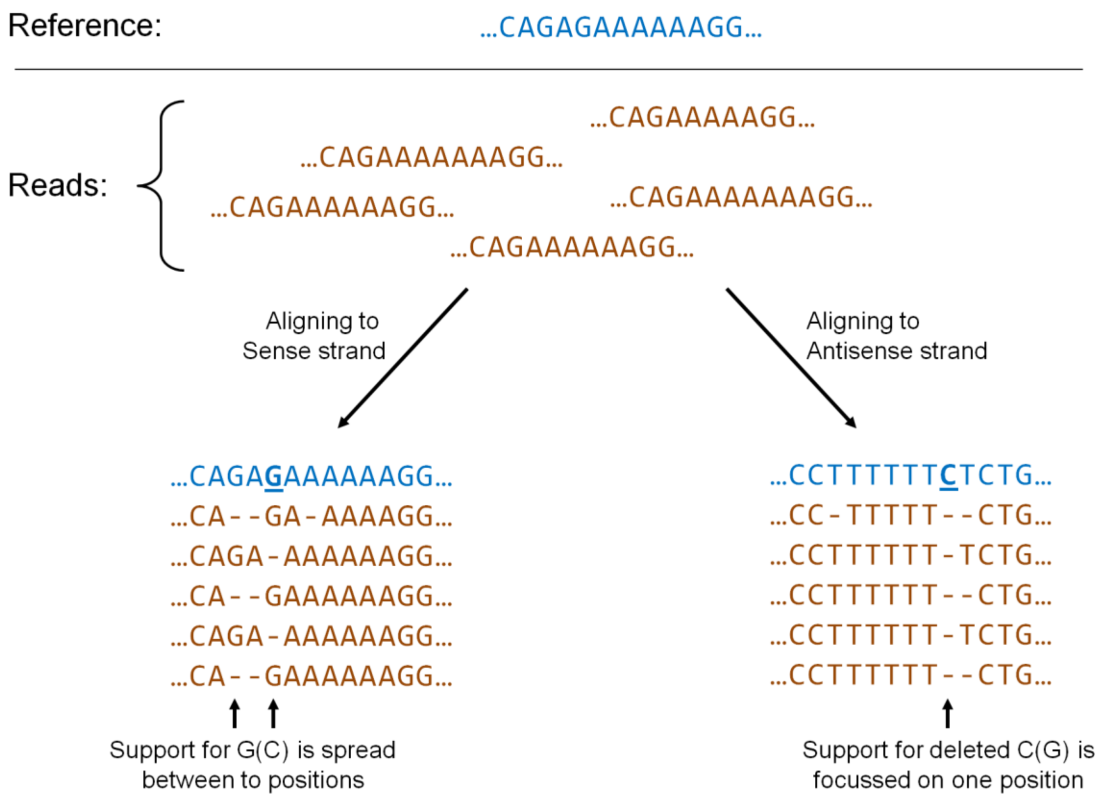


**Figure S2. An illustrative example of inhibited variant detection near a homopolymer due to an artifact of left-aligning indels.** The reference sequence that reads are being aligned to is shown in blue. All reads (brown) are missing the "G" base at position 5 of the reference (supporting a deletion), and some reads are also missing "A" bases of the homopolymer (a common error in Nanopore reads). When aligning these reads to the reference sequence as provided (sense strand), due to left-aligning indels, the read support for the "G" base at position 3 gets spread across positions 3 and 5, resulting in the deletion not being reported by the pipeline (bottom left). If the same reads are aligned to the reverse compliment of the reference (antisense strand), left-aligning the indels allows the gaps to fall at the position of the deleted base, resulting in the deletion being detected and reported (bottom right).

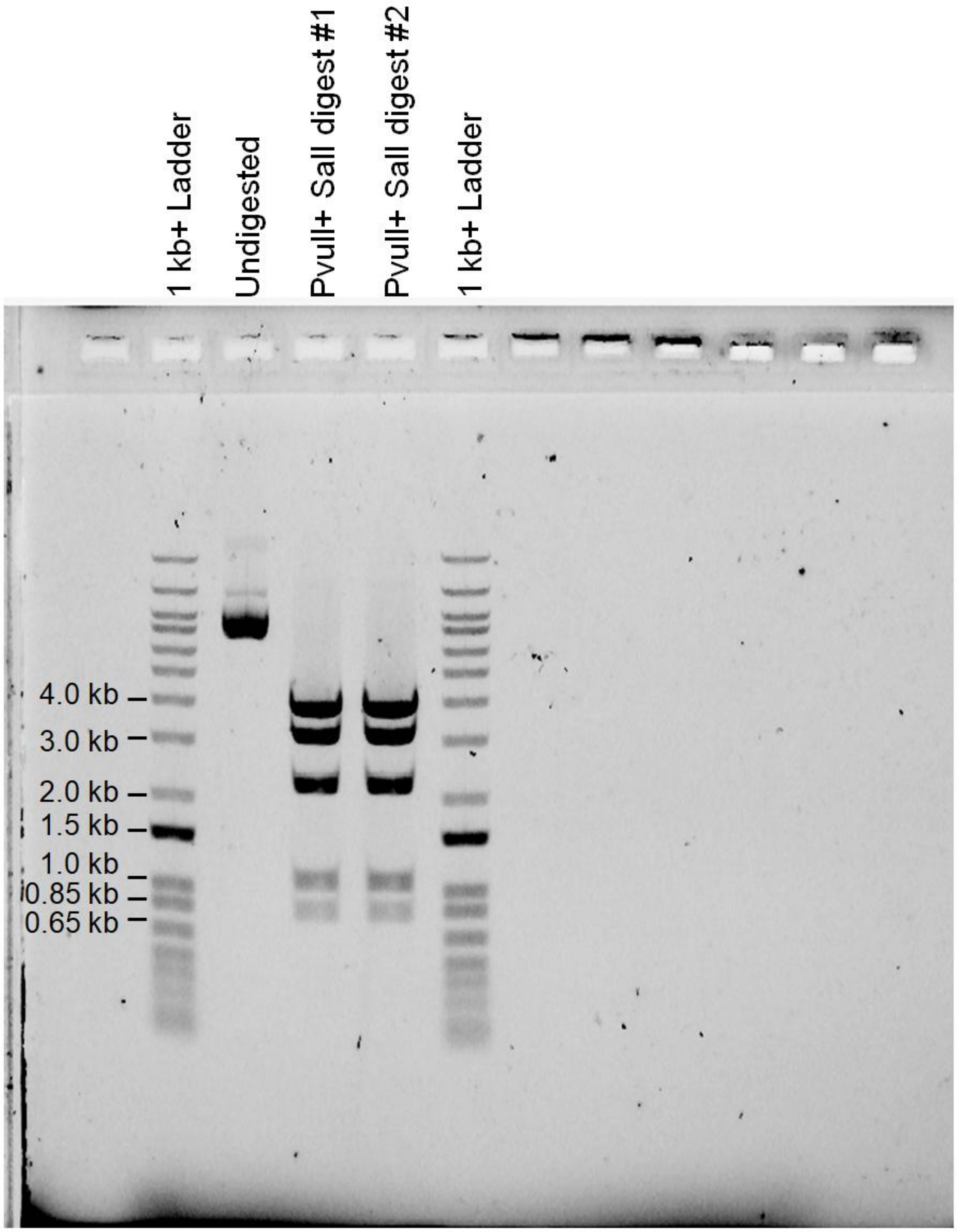


**Figure S3. Representative plasmid fingerprinting gel.** BCRxV.GagPolRev.1 was digested with PvuII-HF and SalI-HF to generate an expected pattern of 3.744, 3.047, 2.154, 1.044 and 0.804 kb bands.

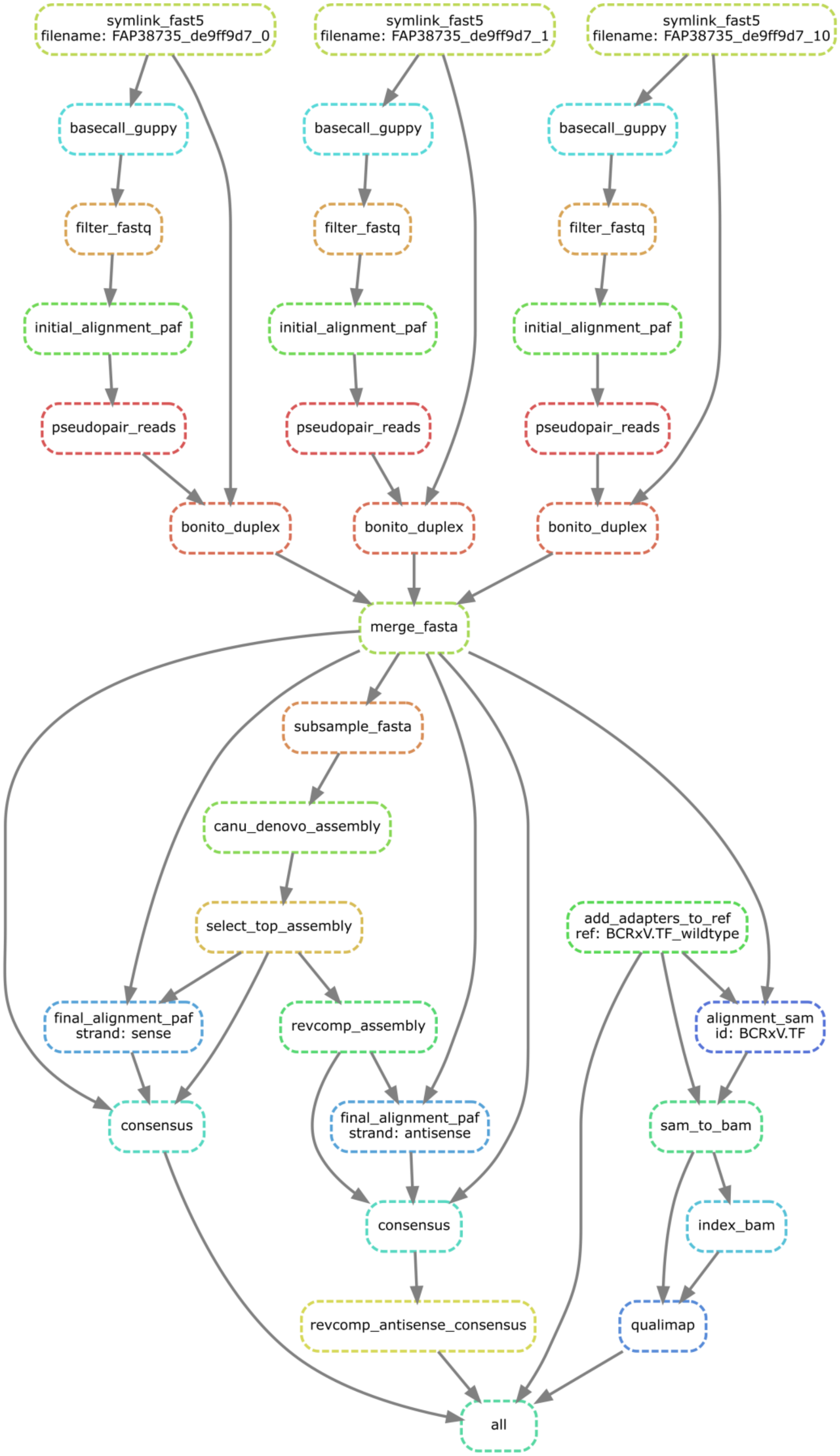


**Figure S4. Flow diagram of Snakemake analysis pipeline.** An overview of the Snakemake pipeline, showing how the inputs and outputs of different rules are connected. The top half of the plot (above “merge_fasta”) is showing the flow for three raw .fast5 files.

**Additional File 1. Reference sequences for the four plasmids sequenced in .fasta format.**
