## Additional File 3 for "Complete sequence verification of plasmid DNA using the Oxford Nanopore Technologies’ MinION device": config.yaml

```
## Reference Sequence
PLASMID_NAME: "BCRxV.VSVG_wildtype"  ## name of reference used for variant detection
REF_SEQ: "/path/to/BCRxV.VSVG_wildtype.fasta"
## allow basecalling to be done using a different reference sequence, used for in silico mutation testing to avoid repeated basecalling for each variant ref.
## when running this pipeline to verify plasmid sequence, this should be the same as the above.
SEQUENCING_REF_SEQ: "/path/to/BCRxV.VSVG_wildtype.fasta"

## Run IDs
CONSENSUS_RUN_ID: "BCRxV.VSVG"

## Storage locations
RESULT_DIR: "/path/to/result_dir"
SCRATCH_DIR: "/path/to/scratch_dir"
FAST5_DIR: "/path/to/raw/minion_data/fast5"

## Cluster headnodes
CPU_CLUSTER: "name_of_cpu_cluster"
CPU_PARTITION: "name_of_cpu_cluster_partition"
GPU_CLUSTER: "name_of_gpu_cluster"
GPU_PARTITION: "name_of_gpu_cluster_partition"

## Guppy basecaller
PATH_TO_GUPPY_INSTALL: "/path/to/bin/ont-guppy_5.0.11/"
GUPPY_MIN_QUAL: 12
## Guppy configs relative to PATH_TO_GUPPY_INSTALL
GUPPY_CONFIG: "data/dna_r10.3_450bps_hac.cfg"
GUPPY_MODEL: "data/template_r10.3_450bps_hac.jsn"

## Bonito basecaller
BONITO_MODEL: "/path/to/minion-plasmid-consensus/data/dna_r10.3_BCRxV.TF_finetuned"

## Sequencing adapters
MINION_ADAPTER_SEQ_5PRIME: "GTTACGTATTGCT"
MINION_ADAPTER_SEQ_3PRIME: "AGCAATAC"

## Parameters - changing these may have unexpected outcomes
READ_LENGTH_WIGGLE: 250            ## +/- number of bases to allow reads to be, compared to ref_seq length.
MIN_BASE_FACTOR: 0.1               ## Minimum depth required at any base to make a call is max(MIN_BASE_DEPTH, average_depth * MIN_BASE_FACTOR).
GLOBAL_THRESHOLD_FACTOR: 5         ## Minimum ratio of called based to second most frequent base (outside of homopolymers)
```
