## Additional File 3 for "Complete sequence verification of plasmid DNA using the Oxford Nanopore Technologies’ MinION device": bonito_duplex_conda_env.yaml

```
name: bonito_duplex
channels:
    - pytorch
dependencies:
    - python==3.7
    - pytorch==1.8.1
    - cudatoolkit=11.1
    - pip:
        - "--editable=git+https://github.com/scottdbrown/seqdist.git@573253e47cbbda3b3cbae02ddc8706fd756e54d0#egg=seqdist"
        - "--editable=git+https://github.com/scottdbrown/bonito.git@23030dd86a232a1bc58012f57ae2f58588393529#egg=ont-bonito"
```
