## Additional File 3 for "Complete sequence verification of plasmid DNA using the Oxford Nanopore Technologies’ MinION device": conda_env.yaml

```
name: MinION_SeqVal_Pipeline
channels:
  - bioconda
  - conda-forge
  - defaults
  - anaconda
dependencies:
  - python==3.7
  - minimap2==2.17
  - samtools==1.7
  - qualimap==2.2.2d
  - biopython==1.77
  
```
