## Additional File 3 for "Complete sequence verification of plasmid DNA using the Oxford Nanopore Technologies’ MinION device": README.md

# MinION Plasmid Sequence Verification pipeline

June 8, 2022

Scott Brown

Citation: TBD

## Purpose

This pipeline is designed to verify the sequence of a plasmid using data from a Oxford Nanopore Technologies' (ONT) MinION sequencing run (flowcell version R10.3).

## Overview

This pipeline is built in snakemake. The flowchart of the pipeline is shown here, shown for three input fast5 files:

<img src="minion_seqval_211028.dag.svg" alt="MinION SeqVal Pipeline"/>

At a high level, this pipeline takes raw `.fast5` files from the MinION, basecalls them using Guppy, filters the resulting passed (Q > 12) `.fastq` on length, aligns the reads to the plasmid reference sequence, pseudopairs forward and reverse reads, uses this pseudopairing as input into Bonito to perform duplex consensus basecalling on the `.fast5` files, and merges the resulting `.fasta` files. 

Next, a subset (500 reads) of this merged `.fasta` is fed into `canu` to create a *de novo* assembly of the reads. All reads in the `.fasta` are aligned to this assembly (sense and antisense versions of the assembly separately to get both right- and left-alignment of potential indels). These alignments are parsed to get a "pileup" at each position. This process selects the most frequent base at each position as the consensus base, and using that generates a consensus sequence for the plasmid. 

As a final step, the user can compare these consensus sequences to the expected reference sequence, confirming that the consensus matches the reference sequence.

Some specific nuances:

* The plasmid reference sequence should be linearized at the site that the digestion occurred during library preparation, and the 5’ overhang should be filled in on both ends. For example, linearizing with BamHI (`G^GATCC`) would result in the linearized reference sequence beginning with `GATCC...` and ending with `...GGATC`.
* This pipeline adds part of the adapter sequence (in lower case bases) to the 5' and 3' end of the plasmid reference sequence (in upper case bases), as this sequence is captured in the sequencing data. This allows the user to compare sequence upstream and downstream of the plasmid sequence. Note that coverage drops in these adapter regions, and it is possible for incorrect bases to be called in these regions without compromising the verification of the plasmid sequence.
* The pipeline has a separate arm that generates a `qualimap` alignment report for the sense strand assembly only. 
* One of the outputs is data to plot a "chromatogram" of the sequence data. This gives the number of reads supporting the top and second top base at each position, and can be plotted in R to visualize the strength of the signal and background noise.
* A global threshold has been added to call the most frequent base, the `GLOBAL_THRESHOLD_FACTOR` - this is set to 5, and is the minimum ratio of top-observed base to second-top-observed base.
* For any position to be called, it must meet a minimum depth set as `max_depth × MIN_BASE_FACTOR`. This ensures that as sequencing depth increases, the boundary between signal and noise is increased proportionally. Note that `MIN_BASE_FACTOR` has a value of 0.1, and the minimum coverage for the plasmid has a value of 100,000, so all bases will have at least 10,000× coverage.
* To get an estimate of the confidence of the consensus sequence, at the consensus generation step, an accuracies file is written which contains the estimated consensus base accuracy at each position of the consensus. This can be used to get a median per-base accuracy across the consensus.

## Requirements

#### General

This pipeline is managed by a `snakemake` workflow, and uses `conda` to manage the packages required. Please make sure you have the following software installed, version numbers represent the version that has been tested to work, but other versions may also work:

* Python - v3.7
* Snakemake - v5.7.4
* conda - v4.8.3

#### Guppy

ONT's Guppy GPU basecaller needs to be installed. This pipeline has been tested with version 5.0.11 [https://mirror.oxfordnanoportal.com/software/analysis/ont-guppy_5.0.11_linux64.tar.gz](https://mirror.oxfordnanoportal.com/software/analysis/ont-guppy_5.0.11_linux64.tar.gz). Copy the tar file to your own directory and unpack as follows:

```bash
(base) [user@server /path/to/bin/Guppy_GPU_v5.0.11]
$ tar -zxvf ont-guppy_5.0.11_linux64.tar.gz
```

#### GSC Servers

This pipeline is designed to be run on BCGSC compute infrastructure, consisting of general purpose linux servers called `gphosts`running `CentOS 7.6`, a CPU cluster (headnode called `numbers`, which uses the `SLURM` scheduler), and a GPU cluster (headnode called `slurmgpu01`, which uses the `SLURM` scheduler) running `CUDA v11` . Modifications will be required to run at your own institution.

Please make sure your user account has passwordless ssh access to both `numbers` and `slurmgpu01`.

#### Fine-tuned Bonito basecalling model

The finetuned Bonito basecalling model is available at [dna_r10.3_BCRxV.TF_finetuned | Zenodo](https://zenodo.org/record/6626041)

[![DOI](https://zenodo.org/badge/DOI/10.5281/zenodo.6626041.svg)](https://doi.org/10.5281/zenodo.6626041)

Download these files and update the path to the Bonito model in the config.yaml file.

## Procedure Work Instructions

#### 1. Copy the raw fast5 data

First we need to copy the raw fast5 sequence data from the Data drive (on the computer connected to the MinION device) to your own networked working directory (if you will not be performing the analysis on the same computer). Copy the entire directory of your sequencing run to a space on your mapped network drive. After the copy is complete, confirm that the directory sizes and number of files are the same on the MinION computer and your working directory.

#### 2. Set up `config.yaml` file

This file sets all the variables that are required for the run. An example `config.yaml` file is included in the root of the repository. The following table explains each variable:

| variable                    | example value                                                         | description                                                                                                                                                                                                                                                                                                                  |
| --------------------------- | --------------------------------------------------------------------- | ---------------------------------------------------------------------------------------------------------------------------------------------------------------------------------------------------------------------------------------------------------------------------------------------------------------------------- |
| `PLASMID_NAME`              | "BCRxV.TF_wildtype"                                                   | Name of plasmid being sequenced.                                                                                                                                                                                                                                                                                             |
| `REF_SEQ`                   | "/path/to/BCRxV.TF.fasta"                                             | Reference sequence of plasmid, linearized at digestion site and with 5' bases filled in as described above.                                                                                                                                                                                                                  |
| `SEQUENCING_REF_SEQ`        | "/path/to/BCRxV.TF.fasta"                                             | Reference sequence of plasmid used for defining forward and reverse reads before Bonito basecalling. This allows multiple variant detection runs to be performed on the same basecalled data, and should only not match the above REF_SEQ during the validation process when testing the detection of *in silico* mutations. |
| `CONSENSUS_RUN_ID`          | "BCRxV.TFg"                                                           | ID for the consensus run.                                                                                                                                                                                                                                                                                                    |
| `RESULT_DIR`                | "/results/dir/"                                                       | Root directory to write the results to. A `CONSENSUS_RUN_ID` folder will be made in this directory.                                                                                                                                                                                                                          |
| `SCRATCH_DIR`               | "/scratch/dir/"                                                       | Root directory for the large interim files (fastq/fasta from basecalling, alignment files). A `CONSENSUS_RUN_ID` folder will be made in this directory.                                                                                                                                                                      |
| `FAST5_DIR`                 | "/path/to/flowcell/fast5/"                                            | Directory containing the raw .fast5 files from the MinION run.                                                                                                                                                                                                                                                               |
| `CPU_CLUSTER`               | "numbers"                                                             | Head node to submit jobs to the CPU cluster.                                                                                                                                                                                                                                                                                 |
| `CPU_PARTITION`             | "upgrade"                                                             | Partition to submit jobs to.                                                                                                                                                                                                                                                                                                 |
| `GPU_CLUSTER`               | "slurmgpu01"                                                          | Head node to submit jobs to the GPU cluster.                                                                                                                                                                                                                                                                                 |
| `GPU_PARTITION`             | "gpu3090"                                                             | Partition to submit jobs to. This must have CUDA v11.                                                                                                                                                                                                                                                                        |
| `PATH_TO_GUPPY_INSTALL`     | "/bin/Guppy_v5.0.11/ont-guppy/"                                       | Root directory of Guppy.                                                                                                                                                                                                                                                                                                     |
| `GUPPY_MIN_QUAL`            | 12                                                                    | Minimum read Q values to set it as `pass` and use in downstream analysis.                                                                                                                                                                                                                                                    |
| `GUPPY_CONFIG`              | "data/dna_r10.3_450bps_hac.cfg"                                       | Guppy config file, relative to `PATH_TO_GUPPY_INSTALL`.                                                                                                                                                                                                                                                                      |
| `GUPPY_MODEL`               | "data/template_r10.3_450bps_hac.jsn"                                  | Guppy model file, relative to `PATH_TO_GUPPY_INSTALL`.                                                                                                                                                                                                                                                                       |
| `BONITO_MODEL`              | "/path/to/minion-plasmid-consensus/data/dna_r10.3_BCRxV.TF_finetuned" | Bonito basecalling model to use.                                                                                                                                                                                                                                                                                             |
| `MINION_ADAPTER_SEQ_5PRIME` | "GTTACGTATTGCT"                                                       | Sequence of 5' adapter that gets added. Example from Ligation Sequencing Kit (SQK-LSK110).                                                                                                                                                                                                                                   |
| `MINION_ADAPTER_SEQ_3PRIME` | "AGCAATAC"                                                            | Sequence of 3' adapter that gets added. Example from Ligation Sequencing Kit (SQK-LSK110).                                                                                                                                                                                                                                   |
| `READ_LENGTH_WIGGLE`        | 250                                                                   | Reads will be filtered to the length of the `SEQUENCING_REF_SEQ` +/- `READ_LENGTH_WIGGLE`.                                                                                                                                                                                                                                   |
| `MIN_BASE_FACTOR`           | 0.1                                                                   | This times the max depth of coverage of the plasmid is the depth-adjusted minimum depth to call a base.                                                                                                                                                                                                                      |
| `GLOBAL_THRESHOLD_FACTOR`   | 5                                                                     | Minimum ratio of called base to second most frequent base.                                                                                                                                                                                                                                                                   |

The following folder structure and files will be created in each of `RESULT_DIR` and `SCRATCH_DIR`:

```
RESULT_DIR:
[CONSENSUS_RUN_ID]
├── ref/
│   └── [PLASMID_NAME]_adapted_sense.fasta
├── qc/
|   ├── [CONSENSUS_RUN_ID]_report.pdf
│   └── genome_results.txt
├── logs/
|   ├── basecalling/
|   |   ├── [fast5_file_name_1]_basecall_guppy_slurm.%N.%j.err
|   |   ├── [fast5_file_name_1]_basecall_guppy_slurm.%N.%j.out
|   |   ├── [fast5_file_name_1]_basecall_guppy_pass_count.txt
|   |   ├── [fast5_file_name_1]_filter_fastq.log
|   |   ├── [fast5_file_name_1]_initial_align_slurm.%N.%j.err
|   |   ├── [fast5_file_name_1]_initial_align_slurm.%N.%j.out
|   |   ├── [fast5_file_name_1]_pseudopair_slurm.%N.%j.err
|   |   ├── [fast5_file_name_1]_pseudopair_slurm.%N.%j.out
|   |   ├── [fast5_file_name_1]_pseudopair_count.txt
|   |   ├── [fast5_file_name_1]_bonito_slurm.%N.%j.err
|   |   ├── [fast5_file_name_1]_bonito_slurm.%N.%j.out
|   |   ├── [fast5_file_name_2]_basecall_guppy_slurm.%N.%j.err
|   |   ...
|   |   └── [fast5_file_name_2]_bonito_slurm.%N.%j.out
|   ├── merge_fasta.log
|   ├── merged_fasta_count.txt
|   ├── [PLASMID_NAME]_add_adapters_to_ref.log
|   ├── canu_assembly_slurm.%N.%j.err
|   ├── canu_assembly_slurm.%N.%j.out
|   ├── select_top_assembly.log
|   ├── revcomp_assembly.log
|   ├── final_align_sense_slurm.%N.%j.err
|   ├── final_align_sense_slurm.%N.%j.out
|   ├── final_align_antisense_slurm.%N.%j.err
|   ├── final_align_antisense_slurm.%N.%j.out
|   ├── consensus_sense_slurm.%N.%j.err
|   ├── consensus_sense_slurm.%N.%j.out
|   ├── consensus_antisense_slurm.%N.%j.err
|   ├── consensus_antisense_slurm.%N.%j.out
|   ├── revcomp_antisense_consensus.log
|   ├── alignment_sam_slurm.%N.%j.err
|   ├── alignment_sam_slurm.%N.%j.out
|   ├── sam_to_bam_slurm.%N.%j.err
|   ├── sam_to_bam_slurm.%N.%j.out
|   ├── index_bam_slurm.%N.%j.err
|   ├── index_bam_slurm.%N.%j.out
|   ├── qualimap_slurm.%N.%j.err
|   └── qualimap_slurm.%N.%j.out
└── consensus/
    ├── sense_chromatogram-data.tsv
    ├── antisense_chromatogram-data.tsv
    ├── sense_consensus.fasta
    ├── sense_per-base-accuracies.tsv
    ├── antisense_consensus.fasta
    ├── antisense_per-base-accuracies.tsv
    └── antisense_consensus_revcomp.fasta
```

```
SCRATCH_DIR:
[CONSENSUS_RUN_ID]
├── 01_fast5/
│   ├── [fast5_file_name_1]
│   │   └── [fast5_file_name_1]             * symlink
│   ├── [fast5_file_name_2]
│   │   └── [fast5_file_name_2]             * symlink
│   ...
├── 02_single_basecalls/
│   ├── [fast5_file_name_1].fastq           * temp file
│   ├── [fast5_file_name_2].fastq           * temp file
│   ...
├── 03_filtered_fastq/
│   ├── [fast5_file_name_1]_filtered.fastq  * temp file
│   ├── [fast5_file_name_2]_filtered.fastq  * temp file
│   ...
├── 04_basecalling_alignment/
│   ├── [fast5_file_name_1].paf             * temp file
│   ├── [fast5_file_name_2].paf             * temp file
│   ...
├── 05_pseudopairs/
│   ├── [fast5_file_name_1].pseudopairs     * temp file
│   ├── [fast5_file_name_2].pseudopairs     * temp file
│   ...
├── 06_paired_basecalls/
│   ├── [fast5_file_name_1].fasta           * temp file
│   ├── [fast5_file_name_1]_summary.tsv     * temp file
│   ├── [fast5_file_name_2].fasta           * temp file
│   ├── [fast5_file_name_2]_summary.tsv     * temp file
│   ...
├── 07_merged_fasta/
|   └── merged.fasta
├── 08_assembly/
│   ├── canu/
│   ├── assembled_sense.fasta
│   └── assembled_antisense.fasta           * temp file
├── 09_alignment/
│   ├── sense_final.paf                     * temp file
│   └── assembled_sense.fasta               * temp file
└── 12_qc_alignment/
    ├── [CONSENSUS_RUN_ID].sam              * temp file
    ├── [CONSENSUS_RUN_ID].bam              * temp file
    └── [CONSENSUS_RUN_ID].bam.bai          * temp file
```

#### 3. Run the `snakemake` workflow

Navigate to the `pipeline` root dir:

```bash
(base) [user@gphost00 ~]
$ cd /path/to/minion-plasmid-consensus/
```

Run the pipeline. Ideally set `-j` to at least 32. Make sure to include the `--use-conda` flag. Override the template configfile with the `--configfile` argument. To prevent apparent missing files due to filesystem latency between gphost and cluster, use the `--latency-wait 60` arg. Add the rule-specific cluster submission arg `--cluster "ssh [USERNAME]@{params.cluster_head_node} 'sbatch {params.cluster_specific_args}'"`, replacing `[USERNAME]` with your username:

```bash
(base) [user@gphost00 /path/to/minion-plasmid-consensus]
$ snakemake -p -j 32 --use-conda --configfile /path/to/config.yaml --latency-wait 60 --cluster "ssh user@{params.cluster_head_node} 'sbatch {params.cluster_specific_args}'"
```

Note that the first time this is run, `snakemake` needs to download and install the conda environment, which may take some time (tests in February 2021 on `gphost08` took 2-3 days. This may be sped up by using [mamba]([GitHub - mamba-org/mamba: The Fast Cross-Platform Package Manager](https://github.com/mamba-org/mamba)).

Basecalling is expected to take up to 5 days (using 8 NVIDIA RTX 3090 GPUs). Alignments and consensus building should then complete within 4-5 hours.

## Check the results

The output of this pipeline is the consensus sequence most-evidenced by the sequence reads (`sense_consensus.fasta` and `antisense_consensus_revcomp.fasta`). These two files are created from alignments to the sense and antisense assemblies, which is done to ensure no variants are missed due to indel alignment biases. Additionally, the pipeline will create an adapted reference sequence with the adapter sequences that flank the linearized plasmid ([PLASMID_NAME]_adapted_sense.fasta).

To verify the plasmid sequence, the user should compare both of the consensus sequences to the adapted reference sequence, and confirm that the sequence between the adapters has a 100% match between the consensus sequence and the plasmid sequence . Note that within the adapters, it is expected that sequence coverage will drop, and mismatches may be present. As long as there does not appear to be a dramatically different sequence in the adapter (indicating a non-adapter sequence present between the plasmid and the adapter), this is acceptable.

It is recommended to do a pairwise alignment of the consensus sequences with the reference sequence to confirm sequence identity between the adapters. Note that the adapter sequences in [PLASMID_NAME]_adapted_sense.fasta are written in lower cases letters to distinguish them from the provided plasmid sequence.

Quality control metrics for the alignment of the reads can been seen in `/RESULT_DIR/CONSENSUS_RUN_ID/qc/CONSENSUS_RUN_ID_report.pdf`.

## Acceptance criteria

* At least 100,000× mean coverage of the plasmid (in the Qualimap QC report).
* All bases of both consensus sequences match the reference sequence exactly for the region between the adapter sequences.
* Median per-base consensus accuracy > 99.9%.
