## Additional File 3 for "Complete sequence verification of plasmid DNA using the Oxford Nanopore Technologies’ MinION device": Snakefile

```
## MinION Sequence verification
## Starting with raw reads and expected reference sequence
## Consensus sequence generation

## Scott Brown

VERSION = "3.1"

## run as:
## snakemake -p -j 32 --use-conda --configfile path/to/config.yaml --latency-wait 60
##    --cluster "ssh user@{params.cluster_head_node} 'sbatch {params.cluster_specific_args}'"


import os
import time
import glob
import logging


###############################
##  Define Global Variables  ##
###############################

configfile: "config.yaml"

## Reference Sequence
REF_NAME = config["PLASMID_NAME"]
REF_SEQ = config["REF_SEQ"]
SEQUENCING_REF_SEQ = config["SEQUENCING_REF_SEQ"]   ## allow basecalling to be done using a different reference
                                                    ## sequence, used for in silico mutation testing to avoid
                                                    ## repeated basecalling for each variant ref.

## Run IDs
CONSENSUS_RUN_ID = config["CONSENSUS_RUN_ID"]

## Storage locations
RESULT_DIR = config["RESULT_DIR"]
SCRATCH_DIR = config["SCRATCH_DIR"]
FAST5_DIR = config["FAST5_DIR"]

## Cluster headnodes
CPU_CLUSTER = config["CPU_CLUSTER"]
CPU_PARTITION = config["CPU_PARTITION"]
GPU_CLUSTER = config["GPU_CLUSTER"]
GPU_PARTITION = config["GPU_PARTITION"]

## Guppy basecaller
PATH_TO_GUPPY_INSTALL = config["PATH_TO_GUPPY_INSTALL"]
GUPPY_MIN_QUAL = config["GUPPY_MIN_QUAL"]
GUPPY_CONFIG = config["GUPPY_CONFIG"]
GUPPY_MODEL = config["GUPPY_MODEL"]

## Bonito basecaller
BONITO_MODEL = config["BONITO_MODEL"]

## Sequencing adapters
MINION_ADAPTER_SEQ_5PRIME = config["MINION_ADAPTER_SEQ_5PRIME"]
MINION_ADAPTER_SEQ_3PRIME = config["MINION_ADAPTER_SEQ_3PRIME"]

## Parameters
READ_LENGTH_WIGGLE = config["READ_LENGTH_WIGGLE"]                       ## +/- number of bases to allow reads to be, compared to REF_LEN.
MIN_BASE_FACTOR = config["MIN_BASE_FACTOR"]                             ## Minimum depth required of any base to make a call is max_depth * MIN_BASE_FACTOR
GLOBAL_THRESHOLD_FACTOR = config["GLOBAL_THRESHOLD_FACTOR"]             ## Minimum ratio of called based to second most frequent base (outside of homopolymers)

## calculate reference length - for filtering reads based on length
REF_LEN = sum([len(x.rstrip().split(">")[0]) for x in open(SEQUENCING_REF_SEQ, "r")]) + len(MINION_ADAPTER_SEQ_5PRIME) + len(MINION_ADAPTER_SEQ_3PRIME)
## count total sequence length of multiline reference fasta, skipping the >header.


## Glob for the fast5 files
## glob returns: ['/path/to/fast5/FAQ22429_755eb6d4_998.fast5',
##                '/path/to/fast5/FAQ22429_755eb6d4_999.fast5']
## and we only want the filename w/o extension
FILENAMES = [os.path.basename(x).split(".")[0] for x in glob.glob(os.path.join(FAST5_DIR, "*.fast5"))]

## Compliment bases
BASE_COMPLIMENT = {"A": "T", "T": "A", "G": "C", "C": "G", "N": "N"}


## Make log folders
#os.makedirs(os.path.join(RESULT_DIR, VAR_DETECTION_RUN_ID, "logs"), exist_ok = True)
os.makedirs(os.path.join(RESULT_DIR, CONSENSUS_RUN_ID, "logs"), exist_ok = True)
os.makedirs(os.path.join(RESULT_DIR, CONSENSUS_RUN_ID, "logs", "basecalling"), exist_ok = True)

localrules: all, symlink_fast5, filter_fastq, add_adapters_to_ref, merge_fasta, subsample_fasta, select_top_assembly, revcomp_assembly, revcomp_antisense_consensus


################
##  Rule All  ##
################

rule all: 
    input:
        expand(os.path.join(RESULT_DIR, CONSENSUS_RUN_ID, "ref", "{ref}_adapted_sense.fasta"), ref = REF_NAME),
        os.path.join(RESULT_DIR, CONSENSUS_RUN_ID, "consensus", "sense_consensus.fasta"),
        os.path.join(RESULT_DIR, CONSENSUS_RUN_ID, "consensus", "antisense_consensus_revcomp.fasta"),
        expand(os.path.join(RESULT_DIR, CONSENSUS_RUN_ID, "qc", "{id}_report.pdf"), id = CONSENSUS_RUN_ID),


########################
##  01 Prepare fast5  ##
########################

rule symlink_fast5:
    '''Create a symlink of each fast5 file in its own directory'''
    input:
        fast5 = os.path.join(FAST5_DIR, "{filename}.fast5")
    output:
        linked_fast5_dir = directory(os.path.join(SCRATCH_DIR, CONSENSUS_RUN_ID, "01_fast5", "{filename}"))
    shell:
        "mkdir {output.linked_fast5_dir}; cd {output.linked_fast5_dir}; ln -s {input.fast5}"


##############################
##  02 Initial Basecalling  ##
##############################

rule basecall_guppy:
    '''Basecalling: generates .fastq files from the raw .fast5 files from the MinION, and count the number of reads'''
    input:
        fast5_dir = rules.symlink_fast5.output.linked_fast5_dir
    output:
        fastq_dir = temp(directory(os.path.join(SCRATCH_DIR, CONSENSUS_RUN_ID, "02_single_basecalls", "{filename}"))),
        pass_count = os.path.join(RESULT_DIR, CONSENSUS_RUN_ID, "logs","basecalling","{filename}_basecall_guppy_pass_count.txt")
    params:
        guppy = os.path.join(PATH_TO_GUPPY_INSTALL, "bin/guppy_basecaller"),
        cnfg = os.path.join(PATH_TO_GUPPY_INSTALL, GUPPY_CONFIG),
        mdl = os.path.join(PATH_TO_GUPPY_INSTALL, GUPPY_MODEL),
        min_qscore = GUPPY_MIN_QUAL,
        gpu_device = '"cuda:0"',    ## since SLURM will give the job a single GPU.
        cluster_head_node = GPU_CLUSTER,
        cluster_specific_args = "--partition={} --mem=12G --gres=gpu:1 -e {} -o {} --job-name=guppy".format(GPU_PARTITION,
            os.path.join(RESULT_DIR, CONSENSUS_RUN_ID, "logs","basecalling","{filename}_basecall_guppy_slurm.%N.%j.err"),
            os.path.join(RESULT_DIR, CONSENSUS_RUN_ID, "logs","basecalling","{filename}_basecall_guppy_slurm.%N.%j.out"))
    shell:
        """{params.guppy} -i {input.fast5_dir} -s {output.fastq_dir} -c {params.cnfg} --model {params.mdl} \
        --min_qscore {params.min_qscore} --device {params.gpu_device}; 
        cat {output.fastq_dir}/pass/*.fastq | echo $((`wc -l`/4)) > {output.pass_count}"""


#########################
##  03 Read Filtering  ##
#########################

rule filter_fastq:
    '''Removes sequences which are too short or too long (not useful)'''
    input:
        fastq_dir = rules.basecall_guppy.output.fastq_dir
    output:
        fastq = temp(os.path.join(SCRATCH_DIR, CONSENSUS_RUN_ID, "03_filtered_fastq", "{filename}_filtered.fastq")),
    params:
        min_length = REF_LEN - READ_LENGTH_WIGGLE,
        max_length = REF_LEN + READ_LENGTH_WIGGLE,
    log: 
        log = os.path.join(RESULT_DIR, CONSENSUS_RUN_ID, "logs","basecalling", "{filename}_filter_fastq.log")
    run:
        logging.basicConfig(filename=log.log, level=logging.INFO)
        try:
            def process(lines=None):
                ks = ['name', 'sequence', 'optional', 'quality']
                return {k: v for k, v in zip(ks, lines)}
            
            out = open(output.fastq, "w")
            setcount = 4
            num_seq_passed = 0
            fastq_file = os.listdir(os.path.join(input.fastq_dir,"pass"))[0]
            with open(os.path.join(input.fastq_dir,"pass",fastq_file), 'r') as fh:
                lines = []
                for line in fh:
                    lines.append(line.rstrip())
                    if len(lines) == setcount:
                        record = process(lines)

                        if len(record["sequence"]) >= params.min_length and len(record["sequence"]) <= params.max_length:
                            out.write("{}\n{}\n{}\n{}\n".format(record["name"], record["sequence"], record["optional"], record["quality"]))
                            num_seq_passed += 1
                        lines = []
            out.close()
            logging.info("{}\n".format(num_seq_passed))
        except Exception as e: 
            logging.error(e, exc_info=True)
            raise


############################
##  04 Initial Alignment  ##
############################

rule initial_alignment_paf:
    '''Generate the .paf alignment file for determining forward vs. reverse reads'''
    input:
        fastq = rules.filter_fastq.output.fastq,
        refseq = SEQUENCING_REF_SEQ
    output:
        paf = temp(os.path.join(SCRATCH_DIR, CONSENSUS_RUN_ID, "04_basecalling_alignment", "{filename}.paf"))
    params:
        cluster_head_node = CPU_CLUSTER,
        cluster_specific_args = "--partition={} --mem=2G -e {} -o {} --job-name=initial_minimap2".format(CPU_PARTITION,
            os.path.join(RESULT_DIR, CONSENSUS_RUN_ID, "logs","basecalling", "{filename}_initial_align_slurm.%N.%j.err"),
            os.path.join(RESULT_DIR, CONSENSUS_RUN_ID, "logs","basecalling", "{filename}_initial_align_slurm.%N.%j.out"))
    conda: "env/conda_env.yaml"
    shell:
        "minimap2 -x map-ont --cs {input.refseq} {input.fastq} > {output.paf}"


############################
##  05 Pseudo-pair Reads  ##
############################

rule pseudopair_reads:
    '''Generate read pseudo-pairing for use in Bonito'''
    input:
        paf = rules.initial_alignment_paf.output.paf
    output:
        pseudopairs = temp(os.path.join(SCRATCH_DIR, CONSENSUS_RUN_ID, "05_pseudopairs", "{filename}.pseudopairs")),
        pair_count = os.path.join(RESULT_DIR, CONSENSUS_RUN_ID, "logs","basecalling","{filename}_pseudopair_count.txt")
    params:
        script = "src/pseudopair_reads.py",
        min_align = REF_LEN - (2*READ_LENGTH_WIGGLE),
        cluster_head_node = CPU_CLUSTER,
        cluster_specific_args = "--partition={} --mem=2G -e {} -o {} --job-name=pseudopair".format(CPU_PARTITION,
            os.path.join(RESULT_DIR, CONSENSUS_RUN_ID, "logs","basecalling", "{filename}_pseudopair_slurm.%N.%j.err"),
            os.path.join(RESULT_DIR, CONSENSUS_RUN_ID, "logs","basecalling", "{filename}_pseudopair_slurm.%N.%j.out"))
    conda: "env/conda_env.yaml"
    shell:
        """python {params.script} --paf {input.paf} --min_align_length {params.min_align} --pseudopairs {output.pseudopairs}; 
        cat {output.pseudopairs} | wc -l > {output.pair_count}"""


#############################
##  06 Bonito Basecalling  ##
#############################

rule bonito_duplex:
    '''Perform basecalling on pseudo-paired reads using the Bonito basecaller in duplex mode'''
    input:
        pairs = rules.pseudopair_reads.output.pseudopairs,
        fast5_dir = rules.symlink_fast5.output.linked_fast5_dir
    output:
        fasta = temp(os.path.join(SCRATCH_DIR, CONSENSUS_RUN_ID, "06_paired_basecalls", "{filename}.fasta")),
        summary = temp(os.path.join(SCRATCH_DIR, CONSENSUS_RUN_ID, "06_paired_basecalls", "{filename}_summary.tsv"))
    params:
        gpu_device = '"cuda:0"',    ## since SLURM will give the job a single GPU.
        max_cpus = 4,
        model = BONITO_MODEL,
        cluster_head_node = GPU_CLUSTER,
        cluster_specific_args = "--partition={} --mem=40000 --gres=gpu:1 -n 6 -e {} -o {} --job-name=bonito".format(GPU_PARTITION,
            os.path.join(RESULT_DIR, CONSENSUS_RUN_ID, "logs","basecalling", "{filename}_bonito_slurm.%N.%j.err"),
            os.path.join(RESULT_DIR, CONSENSUS_RUN_ID, "logs","basecalling", "{filename}_bonito_slurm.%N.%j.out"))
    conda: "env/bonito_duplex_conda_env.yaml"
    shell:
        """
        if [[ $(wc -l <{input.pairs}) -gt 0 ]]
        then
          bonito duplex --pairs {input.pairs} --max-cpus {params.max_cpus} --device {params.gpu_device} \
        {params.model} {input.fast5_dir}/ > {output.fasta}
        else
          touch {output.fasta}; touch {output.summary};
        fi
        """


######################
##  07 Merge Fasta  ##
######################

rule merge_fasta:
    '''Merges all the separate fasta files generated from basecalling'''
    input:
        expand(rules.bonito_duplex.output.fasta, filename = FILENAMES)
    output:
        merged_fasta = os.path.join(SCRATCH_DIR, CONSENSUS_RUN_ID, "07_merged_fasta", "merged.fasta"),
        merged_count = os.path.join(RESULT_DIR, CONSENSUS_RUN_ID, "logs","merged_fasta_count.txt")
    params:
        fasta_dir = os.path.join(SCRATCH_DIR, CONSENSUS_RUN_ID, "06_paired_basecalls")
    log: os.path.join(RESULT_DIR, CONSENSUS_RUN_ID, "logs", "merge_fasta.log")
    shell:
        """cat {params.fasta_dir}/*.fasta > {output.merged_fasta} 2>&1 | tee {log}; 
        grep -c '>' {output.merged_fasta} > {output.merged_count}"""


###########################
##  08 De novo assembly  ##
###########################

rule subsample_fasta:
    '''Creates a subsampled set of the merged fasta to use for draft de novo assembly'''
    input:
        rules.merge_fasta.output.merged_fasta
    output:
        sub_fasta = temp(os.path.join(SCRATCH_DIR, CONSENSUS_RUN_ID, "08_assembly", "subsampled.fasta"))
    params:
        lines = 1000
    log: os.path.join(RESULT_DIR, CONSENSUS_RUN_ID, "logs", "subsample_fasta.log")
    shell:
        "head -{params.lines} {input} > {output.sub_fasta} 2>&1 | tee {log};"
        
rule canu_denovo_assembly:
    '''A draft de novo assembly is done to create a reference to align the reads to when determining consensus'''
    input:
        reads = rules.subsample_fasta.output.sub_fasta
    output:
        unitigs = os.path.join(SCRATCH_DIR, CONSENSUS_RUN_ID, "08_assembly", "canu", "assembled.unitigs.fasta"),
    params:
        outdir = os.path.join(SCRATCH_DIR, CONSENSUS_RUN_ID, "08_assembly", "canu"),
        plasmid_size = REF_LEN,
        cluster_head_node = CPU_CLUSTER,
        cluster_specific_args = "--partition={} --mem 4G -e {} -o {} -n 16 --job-name=canu_assembly".format(CPU_PARTITION,
            os.path.join(RESULT_DIR, CONSENSUS_RUN_ID, "logs", "canu_assembly_slurm.%N.%j.err"),
            os.path.join(RESULT_DIR, CONSENSUS_RUN_ID, "logs", "canu_assembly_slurm.%N.%j.out"))
    conda: "env/conda_env_canu.yaml"
    shell:
        "canu -assemble -d {params.outdir} -p assembled genomeSize={params.plasmid_size} useGrid=false maxThreads=16 -corrected -nanopore {input.reads}"

rule select_top_assembly:
    '''Selects the first fasta entry in input.unitigs, in the case that there is more than one assembly'''
    input:
        unitigs = rules.canu_denovo_assembly.output.unitigs
    output:
        top_assembly = os.path.join(SCRATCH_DIR, CONSENSUS_RUN_ID, "08_assembly", "assembled_sense.fasta")
    log:
        log = os.path.join(RESULT_DIR, CONSENSUS_RUN_ID, "logs", "select_top_assembly.log")
    run:
        logging.basicConfig(filename=log.log, level=logging.INFO)
        try:
            out = open(output.top_assembly, "w")
            seqs = [""]
            i = -1
            for line in open(input.unitigs, "r"):
                if line.startswith(">"):
                    i += 1
                    if len(seqs) < i:
                        seqs.append("")
                    seqs[i] = line
                else:
                    ## save multi-line fasta seq
                    seqs[i] += line.rstrip()
            # write first seq
            out.write("{}\n".format(seqs[0]))
            out.close()
            logging.info("Completed.\n")
        except Exception as e: 
            logging.error(e, exc_info=True)
            raise

rule revcomp_assembly:
    '''Creates a reverse-complemented assembly sequence for increased sensitivity of consensus base calling'''
    input:
        refseq = rules.select_top_assembly.output.top_assembly
    output:
        revcomp_refseq = temp(os.path.join(SCRATCH_DIR, CONSENSUS_RUN_ID, "08_assembly", "assembled_antisense.fasta"))
    log:
        log = os.path.join(RESULT_DIR, CONSENSUS_RUN_ID, "logs", "revcomp_assembly.log")
    run:
        logging.basicConfig(filename=log.log, level=logging.INFO)
        try:
            out = open(output.revcomp_refseq, "w")
            sense_seq = ""
            for line in open(input.refseq, "r"):
                if line.startswith(">"):
                    out.write(line)
                else:
                    ## save multi-line fasta seq
                    sense_seq += line.rstrip()
            #reverse sequence, and for each base write the compliment.
            out.write("".join([BASE_COMPLIMENT[x.upper()] for x in line.rstrip()[::-1]]))
            out.close()
            logging.info("Completed.\n")
        except Exception as e: 
            logging.error(e, exc_info=True)
            raise


##########################
##  09 Final Alignment  ##
##########################

rule final_alignment_paf:
    '''Generate the .paf alignment file for determining consensus sequence'''
    input:
        fasta = rules.merge_fasta.output.merged_fasta,
        refseq = os.path.join(SCRATCH_DIR, CONSENSUS_RUN_ID, "08_assembly", "assembled_{strand}.fasta")
    output:
        paf = temp(os.path.join(SCRATCH_DIR, CONSENSUS_RUN_ID, "09_alignment", "{strand}_final.paf"))
    params:
        cluster_head_node = CPU_CLUSTER,
        cluster_specific_args = "--partition={} --mem 30G -e {} -o {} --job-name=final_minimap2".format(CPU_PARTITION,
            os.path.join(RESULT_DIR, CONSENSUS_RUN_ID, "logs", "final_align_{strand}_slurm.%N.%j.err"),
            os.path.join(RESULT_DIR, CONSENSUS_RUN_ID, "logs", "final_align_{strand}_slurm.%N.%j.out"))
    conda: "env/conda_env.yaml"
    shell:
        "minimap2 -x map-ont --cs {input.refseq} {input.fasta} > {output.paf}"


##################################
##  10 Consensus Determination  ##
##################################

rule consensus:
    '''Pileup base called at each position and generate consensus sequence and "chromatogram" trace file'''
    input:
        paf = rules.final_alignment_paf.output.paf,
        refseq = os.path.join(SCRATCH_DIR, CONSENSUS_RUN_ID, "08_assembly", "assembled_{strand}.fasta"),
        raw_reads = rules.merge_fasta.output.merged_fasta
    output:
        consensus = os.path.join(RESULT_DIR, CONSENSUS_RUN_ID, "consensus", "{strand}_consensus.fasta"),
        chromat = os.path.join(RESULT_DIR, CONSENSUS_RUN_ID, "consensus", "{strand}_chromatogram-data.tsv"),
        accuracies = os.path.join(RESULT_DIR, CONSENSUS_RUN_ID, "consensus", "{strand}_per-base-accuracies.tsv")
    params:
        script = "src/mapped_paf_read_parser.py",
        min_depth_factor = MIN_BASE_FACTOR,
        global_threshold = GLOBAL_THRESHOLD_FACTOR,
        cluster_head_node = CPU_CLUSTER,
        cluster_specific_args = "--partition={} --mem 4G -e {} -o {} --job-name=consensus".format(CPU_PARTITION,
            os.path.join(RESULT_DIR, CONSENSUS_RUN_ID, "logs", "consensus_{strand}_slurm.%N.%j.err"),
            os.path.join(RESULT_DIR, CONSENSUS_RUN_ID, "logs", "consensus_{strand}_slurm.%N.%j.out"))
    conda: "env/conda_env.yaml"
    shell:
        """python {params.script} --ref {input.refseq} --paf {input.paf} --reads {input.raw_reads} \
        --consensus {output.consensus} --chromat {output.chromat} --accuracies {output.accuracies} \
        --min_depth_factor {params.min_depth_factor} --global_threshold_factor {params.global_threshold} """

rule revcomp_antisense_consensus:
    '''Creates a reverse-complemented consensus sequence for ease of comparison to expected reference'''
    input:
        antisense_consensus = os.path.join(RESULT_DIR, CONSENSUS_RUN_ID, "consensus", "antisense_consensus.fasta")
    output:
        revcomp_consensus = os.path.join(RESULT_DIR, CONSENSUS_RUN_ID, "consensus", "antisense_consensus_revcomp.fasta")
    log:
        log = os.path.join(RESULT_DIR, CONSENSUS_RUN_ID, "logs", "revcomp_antisense_consensus.log")
    run:
        logging.basicConfig(filename=log.log, level=logging.INFO)
        try:
            out = open(output.revcomp_consensus, "w")
            sense_seq = ""
            for line in open(input.antisense_consensus, "r"):
                if line.startswith(">"):
                    out.write(line)
                else:
                    ## save multi-line fasta seq
                    sense_seq += line.rstrip()
            #reverse sequence, and for each base write the compliment.
            out.write("".join([BASE_COMPLIMENT[x.upper()] for x in line.rstrip()[::-1]]))
            out.close()
            logging.info("Completed.\n")
        except Exception as e: 
            logging.error(e, exc_info=True)
            raise


###################################
##  11 Expected Reference Prep.  ##
###################################

rule add_adapters_to_ref:
    '''To aid in alignment and consensus base calling near ends of template, MinION adapter sequences 
       (that are captured by sequencing) are added to beginning and end of template'''
    input:
        refseq = REF_SEQ
    output:
        adapted_refseq = os.path.join(RESULT_DIR, CONSENSUS_RUN_ID, "ref", "{ref}_adapted_sense.fasta"),
    params:
        ADAPTER_SEQ_5PRIME = MINION_ADAPTER_SEQ_5PRIME,
        ADAPTER_SEQ_3PRIME = MINION_ADAPTER_SEQ_3PRIME,
    log:
        log = os.path.join(RESULT_DIR, CONSENSUS_RUN_ID, "logs", "{ref}_add_adapters_to_ref.log")
    run:
        logging.basicConfig(filename=log.log, level=logging.INFO)
        try:
            out = open(output.adapted_refseq, "w")
            WRITTEN_5PRIME = False
            for line in open(input.refseq, "r"):
                if line.startswith(">"):
                    out.write(line)
                elif not WRITTEN_5PRIME:
                    out.write(params.ADAPTER_SEQ_5PRIME.lower())
                    WRITTEN_5PRIME = True
                    out.write(line.rstrip().upper())
                else:
                    out.write(line.rstrip().upper())
            ## write 3'
            out.write(params.ADAPTER_SEQ_3PRIME.lower())
            out.close()
            logging.info("Completed.\n")
        except Exception as e: 
            logging.error(e, exc_info=True)
            raise


#############
##  12 QC  ##
#############

rule alignment_sam:
    '''Generate the .sam alignment file for QC report'''
    input:
        fasta = rules.merge_fasta.output.merged_fasta,
        #refseq = os.path.join(SCRATCH_DIR, CONSENSUS_RUN_ID, "08_assembly", "assembled_sense.fasta"),
        refseq = expand(rules.add_adapters_to_ref.output.adapted_refseq, ref = REF_NAME)
    output:
        sam = temp(os.path.join(SCRATCH_DIR, CONSENSUS_RUN_ID, "12_qc_alignment", "{id}.sam"))
    params:
        cluster_head_node = CPU_CLUSTER,
        cluster_specific_args = "--partition={} --mem 30G -e {} -o {} --job-name=sam_minimap2".format(CPU_PARTITION,
            os.path.join(RESULT_DIR, CONSENSUS_RUN_ID, "logs", "alignment_sam_slurm.%N.%j.err"),
            os.path.join(RESULT_DIR, CONSENSUS_RUN_ID, "logs", "alignment_sam_slurm.%N.%j.out"))
    conda: "env/conda_env.yaml"
    shell:
        "minimap2 -ax map-ont --cs {input.refseq} {input.fasta} > {output.sam}"

rule sam_to_bam:
    '''Convert .sam to .bam for QC report'''
    input:
        sam = rules.alignment_sam.output.sam,
        #refseq = os.path.join(SCRATCH_DIR, CONSENSUS_RUN_ID, "08_assembly", "assembled_sense.fasta"),
        refseq = expand(rules.add_adapters_to_ref.output.adapted_refseq, ref = REF_NAME)
    output:
        bam = temp(os.path.join(SCRATCH_DIR, CONSENSUS_RUN_ID, "12_qc_alignment", "{id}.bam"))
    params:
        cluster_head_node = CPU_CLUSTER,
        cluster_specific_args = "--partition={} --mem 30G -e {} -o {} --job-name=sambam_samtools".format(CPU_PARTITION,
            os.path.join(RESULT_DIR, CONSENSUS_RUN_ID, "logs", "sam_to_bam_slurm.%N.%j.err"),
            os.path.join(RESULT_DIR, CONSENSUS_RUN_ID, "logs", "sam_to_bam_slurm.%N.%j.out"))
    conda: "env/conda_env.yaml"
    shell:
        "samtools view -bT {input.refseq} {input.sam} | samtools sort -o {output.bam}"

rule index_bam:
    '''Index bam for QC report'''
    input:
        bam = rules.sam_to_bam.output.bam
    output:
        bai = temp(os.path.join(SCRATCH_DIR, CONSENSUS_RUN_ID, "12_qc_alignment", "{id}.bam.bai"))
    params:
        cluster_head_node = CPU_CLUSTER,
        cluster_specific_args = "--partition={} --mem 30G -e {} -o {} --job-name=index_samtools".format(CPU_PARTITION,
            os.path.join(RESULT_DIR, CONSENSUS_RUN_ID, "logs", "index_bam_slurm.%N.%j.err"),
            os.path.join(RESULT_DIR, CONSENSUS_RUN_ID, "logs", "index_bam_slurm.%N.%j.out"))
    conda: "env/conda_env.yaml"
    shell:
        "samtools index {input.bam}"

rule qualimap:
    '''Generate QC report of alignment using qualimap'''
    input:
        bam = rules.sam_to_bam.output.bam,
        bai = rules.index_bam.output.bai
    output:
        pdf = os.path.join(RESULT_DIR, CONSENSUS_RUN_ID, "qc", "{id}_report.pdf")
    params:
        outdir = os.path.join(RESULT_DIR, CONSENSUS_RUN_ID, "qc"),
        outfile = lambda wildcards: "{}_report.pdf".format(wildcards.id),
        outformat = "PDF",
        java_mem = "16G",
        cluster_head_node = CPU_CLUSTER,
        cluster_specific_args = "--partition={} --mem 30G -n 32 -e {} -o {} --job-name=qualimap".format(CPU_PARTITION,
            os.path.join(RESULT_DIR, CONSENSUS_RUN_ID, "logs", "qualimap_slurm.%N.%j.err"),
            os.path.join(RESULT_DIR, CONSENSUS_RUN_ID, "logs", "qualimap_slurm.%N.%j.out"))
    conda: "env/conda_env.yaml"
    threads: 32
    shell:
        """qualimap bamqc -nt {threads} --java-mem-size={params.java_mem} -bam {input.bam} -outdir {params.outdir} \
        -outformat {params.outformat} -outfile {params.outfile}"""
```
