## Additional File 3 for "Complete sequence verification of plasmid DNA using the Oxford Nanopore Technologies’ MinION device": mapped_paf_read_parser.py

```
TITLE = "Mapped PAF Read Parser"
DESC = "Parse the minimap2 mapped reads, and determine the consensus sequence"
'''
@author: sbrown
'''

VERSION = "5.1"

## Run as:
## python {params.script} --ref {input.refseq} --reads {input.reads} --paf {input.paf} --consensus {output.consensus} --chromat {output.chromat} 
#   --accuracies {output.accuracies} --min_depth_factor {params.mdf} --global_threshold_factor {params.gtf}

## Import Libraries

import sys
import argparse
import os
import time


## Declare global variables

DEBUG = False
VERB = False

## Compliment bases
BASE_COMPLIMENT = {"A": "T", "T": "A", "G": "C", "C": "G", "N": "N"}

## Classes and functions

def statprint(msg, msg_type = "STATUS"):
    print("{message_type} [{datetime}]: {message}".format(message_type = msg_type,
             datetime = time.strftime("%Y/%m/%d %T"), 
             message = msg))


def processBaseString_leftIndel(obsarr, obsarr_i, baseseqstr):
    ## since I want to right-align all the base strings, but i don't know what the longest string will be,
    ## I am inserting new positions to the front of the list.
    ## 
    ## Read 1:  TAAT
    ## Read 2:  AT
    ## 
    ## Right-aligned: TAAT
    ##                  AT
    ##
    ## Stored:
    ##   obsarr[i][0] = {'A': 0, 'T': 1, 'C': 0, 'G': 0}
    ##   obsarr[i][1] = {'A': 1, 'T': 0, 'C': 0, 'G': 0}
    ##   obsarr[i][2] = {'A': 2, 'T': 0, 'C': 0, 'G': 0}
    ##   obsarr[i][3] = {'A': 0, 'T': 2, 'C': 0, 'G': 0}

    ## we iterate through the base string to add, in reverse (right-aligned string, left-aligned indel)
    ## bi will index from 0:len(baseseqstr), and we use that to iterate through bases of baseseqstr from right to left.
    for bi in range(len(baseseqstr)):  ## iterate through positions of bases string
        b = baseseqstr[len(baseseqstr) - bi - 1]  ## get base starting from 3' end
        if len(obsarr[obsarr_i]) <= bi:
            # insert a position at the front of the list - not ideal for efficiency
            obsarr[obsarr_i].insert(0, {'A': 0, 'T': 0, 'C': 0, 'G': 0})
        ## update the bi position (from end of list)
        obsarr[obsarr_i][-(1+bi)][b] += 1
    return obsarr

def processBaseString_rightIndel(obsarr, obsarr_i, baseseqstr):
    ## we iterate through the base string to add (left-aligned string, right-aligned indel)
    ## bi will index through obsarr[obsarr_i] left to right
    for bi in range(len(baseseqstr)):  ## iterate through positions of bases string
        b = baseseqstr[bi]  ## get base starting from 5' end
        if len(obsarr[obsarr_i]) <= bi:
            obsarr[obsarr_i].append({'A': 0, 'T': 0, 'C': 0, 'G': 0})
        obsarr[obsarr_i][bi][b] += 1
    return obsarr

def processOperation(obsarr, i, operator, operand, refarr):
    if operator == ":":
        ## bases match
        for x in range(int(operand)):
            ## x is used just to iterate through the number of matched bases, but i is the index we care about.
            obsarr = processBaseString_leftIndel(obsarr, (2*i)+1, refarr[(2*i)+1])
            i += 1
    elif operator == "+":
        ## insertion of bases
        inserted = ""
        for k in operand:
            ## k == each base that is inserted
            inserted += k
        obsarr = processBaseString_leftIndel(obsarr, 2*i, inserted.upper())
    elif operator == "-":
        ## deletion of bases
        ## don't write anything, does not contribute to depth.
        for x in range(len(operand)):
            i += 1
    elif operator == "*":
        ## substitution of bases
        ## write second base of operand
        obsarr = processBaseString_leftIndel(obsarr, (2*i)+1, operand[-1].upper())
        i += 1
    elif operator == "Z":
        pass
    else:
        ## unknown operator, error.
        sys.exit("Unknown operator: {}".format(operator))
    
    return (obsarr, i)

## Main

if __name__ == "__main__":

    ## Deal with command line arguments
    parser = argparse.ArgumentParser(description = TITLE)
    parser.add_argument("--ref", dest = "REF", help = "Reference fasta file of plasmid sequence", type = str)
    parser.add_argument("--reads", dest = "READS", help = "Raw reads fasta file", type = str)
    parser.add_argument("--paf", dest = "PAF", help = "Mapped reads .paf file", type = str)
    parser.add_argument("--consensus", help = "Consensus output file", type = str)
    parser.add_argument("--chromat", help = "Chromatogram data output file", type = str)
    parser.add_argument("--accuracies", help = "Per position consensus accuracies output file", type = str)
    parser.add_argument("--min_depth_factor", dest = "MIN_DEPTH_FACTOR", help = "Minimum number of reads needed to call a base is set to max_depth*MIN_DEPTH_FACTOR", type = float)
    parser.add_argument("--global_threshold_factor", dest = "GLOBAL_THRESHOLD_FACTOR", help = "Value of minimum most frequent to second most frequent base ratio to make call", type = float)
    parser.add_argument("-d", "--debug", action = "store_true", dest = "DEBUG", help = "Flag for setting debug/test state.")
    parser.add_argument("-v", "--verbose", action = "store_true", dest = "VERB", help = "Flag for setting verbose output.")
    args = parser.parse_args()

    ## Set Global Vars
    DEBUG = args.DEBUG
    VERB = args.VERB

    print("=======================================================")
    print("Python version: {}".format(sys.version))
    print("Python environment: {}".format(sys.prefix))
    print("Server: {}".format(os.uname()[1]))
    print("Current directory: {}".format(os.getcwd()))
    print("Command: {}".format(" ".join(sys.argv)))
    print("Time: {}".format(time.strftime("%Y/%m/%d %T")))
    print("=======================================================\n")
    
    ## Overview:
    ## 1. Initialize an array of the reference sequence
    ## 2. Read in .paf file and extract relevant fields for each alignment
    ## 3. Read in raw reads to extract seq before and after alignments
    ## 4. Parse through each alignment from .paf file and iterate through the cs tag
    ##    Determine the observed nucleotide frequency at each position of the reference
    ## 5. Determine average coverage across template
    ## 6. Determine the consensus sequence based on the most observed nucleotide at each position
    ## 7. Output the consensus and frequency at each position for plotting.


    ########################################
    ## Step 1: Initialize Reference Array ##
    ########################################

    ## Position n of reference sequence will be at position 2n+1 of reference array
    ## Position 2n of reference array will hold additional bases in the case of insertions between position n-1 and n of reference sequence
    '''
    Reference:  A C C A T G
                0 1 2 3 4 5
    Array:     ['', A, '',  C, '',  C, '',  A, '',  T, '',  G, '']
                0   1   2   3   4   5   6   7   8   9  10  11  12
    '''

    statprint("Initializing reference array...")
    refseq = ""
    for line in open(args.REF, "r"):
        if not line.startswith(">"):
            refseq += line.rstrip().upper()
    
    refarr = ["" for x in range((2*len(refseq))+1)] ## to hold the reference sequence
    obsarr = [[] for x in range((2*len(refseq))+1)] ## to hold the actual observed bases

    ## ultimately obsarr will be obsarr[array_position][base_position_from_5'_end]{dict, of, base, counts}
    ## this allows for one array_position to hold more than a single base for each read (ie an insertion of 2 bases)
    ## In cases where there is more than 1 base, we right-align* all the possible bases across the reads (left-aligning indel):
    ## * except for read sequence downstream of an alignment end.
    ##    base pos:    0 1 2
    ##     Read 1:       A T
    ##     Read 2:         T
    ##     Read 3:     C A A
    ## So, obsarr[array_position] = [{C: 1}, {A: 2}, {T: 2, A: 1}]
    ## Then, when checking each position, we step through base_positions at array_position

    ## initialize the reference sequence
    ## This is used to determine the base when an alignment matches the reference.
    for i in range(len(refseq)):
        refarr[(2*i)+1] = refseq[i]


    ################################
    ## Step 2: Read in alignments ##
    ################################

    ## read in paf file
    statprint("Reading in aligments...")
    num_mapped_reads = 0    ## this actually counts alignments, not mapped reads, per se.
    paf = {}

    for line in open(args.PAF, "r"):
        line = line.rstrip().split("\t")
        ## PAF format:
        # Ind   Type        Description
        # 0     string      Query sequence name
        # 1     int         Query sequence length
        # 2     int         Query start coordinate (0-based)
        # 3     int         Query end coordinate (0-based)
        # 4     char        ‘+’ if query/target on the same strand; ‘-’ if opposite
        # 5     string      Target sequence name
        # 6     int         Target sequence length
        # 7     int         Target start coordinate on the original strand
        # 8     int         Target end coordinate on the original strand
        # 9     int         Number of matching bases in the mapping
        # 10    int         Number bases, including gaps, in the mapping
        # 11    int         Mapping quality (0-255 with 255 for missing)

        num_mapped_reads += 1

        if num_mapped_reads % 100000 == 0:
            statprint("{} alignments have been read in.".format(num_mapped_reads))

        read_name = line[0]
        read_length = int(line[1])
        align_start_pos = int(line[2])
        align_end_pos = int(line[3])    ## note this is the index you would use in python subsetting [align_start_pos:align_end_pos]
        ref_start_pos = int(line[7])
        strand = line[4]

        ## reverse indices if reverse alignment
        if strand == "-":
            align_start_pos = read_length - align_end_pos    # 0 based index of reveresed fastq record.
            align_end_pos = read_length - int(line[2])

        ## the position of the cs tag is variable, so step through and find it
        cstag = [e for e in line if e.startswith("cs:")][0]
        
        # cleave off 'cs:'
        cstag = cstag[3:]

        ## save record
        if read_name not in paf:
            ## we only take first alignment from a read
            paf[read_name] = {"align_start_pos": align_start_pos,
                              "align_end_pos": align_end_pos,
                              "ref_start_pos": ref_start_pos,
                              "strand": strand,
                              "cstag": cstag}
    
    statprint("There were {} mapped reads.".format(num_mapped_reads))


    #####################################
    ## Step 3: Read in raw reads fasta ##
    #####################################

    ## this is used to get the read sequence before and after the alignment tracked in the .paf file
    statprint("Reading in fasta file of reads and saving upstream and downstream of alignments...")

    read_name = ""
    seq = ""
    for line in open(args.READS, "r"):
        if line.startswith(">"):
            ## process previous
            if read_name != "" and read_name in paf:
                if paf[read_name]["strand"] == "-":
                    ## revcomp seq
                    seq = "".join([BASE_COMPLIMENT[x.upper()] for x in seq[::-1]])
                paf[read_name]["upstream_seq"] = seq[:paf[read_name]["align_start_pos"]]
                paf[read_name]["downstream_seq"] = seq[paf[read_name]["align_end_pos"]:]
            ## start new fasta entry
            read_name = line.rstrip()[1:]
            seq = ""
        else:
            seq += line.rstrip().upper()
    ## process last entry
    if read_name != "" and read_name in paf:
        if paf[read_name]["strand"] == "-":
            ## revcomp seq
            seq = "".join([BASE_COMPLIMENT[x.upper()] for x in seq[::-1]])
        paf[read_name]["upstream_seq"] = seq[:paf[read_name]["align_start_pos"]]
        paf[read_name]["downstream_seq"] = seq[paf[read_name]["align_end_pos"]:]
    

    ##############################
    ## Step 4: Parse alignments ##
    ##############################

    ## step through alignments and parse the cs tag to determine the matching and mismatching bases
    statprint("Parsing alignments...")

    ## cs tag like:
    ## cs:Z::7-t:112+tt:71-c:49+a:34-ga:3*ct:16*ga:43-t:29+a:1-tta:52+gg:23-a:17+c:2*ag:1-g:198*ca*tc:59
    ## but "cs:" already sliced off the start.
    SPECIAL_CHARS = [":","Z","+","-","*"]
    num_parsed = 0
    for read_name in paf:
        num_parsed += 1

        if num_parsed % 10000 == 0:
            statprint("{} alignments have been parsed and processed.".format(num_parsed))

        operand = ""    ## specifics of variant, or number of matched bases
        operator = ""   ## one of the SPECIAL_CHARS that tells you what action is happening
        i = paf[read_name]["ref_start_pos"]

        ## process prefix of read before it aligns
        obsarr = processBaseString_leftIndel(obsarr, 2*i, paf[read_name]["upstream_seq"])
        
        ## step through and parse
        for c in paf[read_name]["cstag"]:
            if c in SPECIAL_CHARS:
                ## check if existing operator to do
                if operand != "":
                    ## do the previous operator ##
                    (obsarr, i) = processOperation(obsarr, i, operator, operand, refarr)
                
                ## reset the operator
                operator = c
                operand = ""
            else:
                operand += c ## add character to the operator, since it can be multiple characters

        ## end of cstag, do the last operator
        (obsarr, i) = processOperation(obsarr, i, operator, operand, refarr)

        ## process suffix of read after it aligns
        obsarr = processBaseString_rightIndel(obsarr, 2*i, paf[read_name]["downstream_seq"])

        ## end of alignment
    
    
    ####################################
    ## Step 5: Calculate max coverage ##
    ####################################

    statprint("Calculating max depth across sequence...")
    max_depth = 0
    for i in range(len(obsarr)):
        if len(obsarr[i]) > 0:
            if sum(obsarr[i][0].values()) > max_depth: ## take first base position of each array position as this will have the max (reversed, left-aligned indels)
                max_depth = sum(obsarr[i][0].values())
    DEPTH_THRESHOLD = max_depth*args.MIN_DEPTH_FACTOR

    statprint("Max depth is {}.".format(max_depth))
    statprint("DEPTH_THRESHOLD is {}.".format(DEPTH_THRESHOLD))


    ###############################################################
    ## Step 6: Calculate consensus and chromatogram trace values ##
    ###############################################################

    statprint("Calculating consensus...")
    consensus = ""                  ## holds built consensus sequence
    consensus_second = ""           ## holds second most frequent base at each position
    consensus_chromat = ""          ## holds built consensus sequence for chromatogram data
    consensus_second_chromat = ""   ## holds second most frequent base at each position for chromatogram data
    countarr = []                   ## holds depth of the base used at each position
    countarr_second = []            ## holds second most frequent base depth at each position
    accuracies = []                 ## hold accuracies of each position of the consensus

    for x in range(len(obsarr)):    ## x is "position", with odds being 2n+1 of pos of ref, evens being in between.

        if x % 1000 == 0:
            statprint("{} positions have been checked...".format(x))

        ## step through possible multiple bases from 5' to 3' (end of array to beginning)
        for base_pos in range(len(obsarr[x])):

            ## Get the Most Frequent base at this position.
            #summary_x = Counter(obsarr[x][base_pos]).most_common(4)
            ## something like [('G', 323), ('T', 12), ('C', 2)]

            ## obsarr[x][base_pos] is like {'A': 5, 'C': 60, 'T': 14, 'G': 6}
            ## convert to list of tuples with no zeros
            list_of_tuples = [(k, v) for k,v in obsarr[x][base_pos].items() if v > 0]

            ## sort this on the values
            summary_x = sorted(list_of_tuples, key=lambda tup: tup[1])[::-1]
            ## now summary_x is like [('C', 60), ('T', 14), ('G', 6), ('A', 5)]

            if len(summary_x) == 0:
                ## no read support at this position
                base = "X"
                count = 0
            elif len(summary_x) == 1:
                ## only one base observed
                base, count = summary_x[0]
            elif summary_x[0][1] > summary_x[1][1]:     ## this could have been joined with previous elif via OR, but kept separate for readability.
                ## there is a distinct top call
                base, count = summary_x[0]
            else:
                ## there is a tie for top call
                ## get all bases with this count
                tied_x = [tup for tup in summary_x if tup[1] == summary_x[0][1]]
                base = "N"
                count = sum([c[1] for c in tied_x])     ## sum the counts of the tied, it is the count of the N.

            ## Get the Second Most Frequent base at this position.
            if len(summary_x) == 0:
                ## no read support at this position
                base2 = "X"
                count2 = 0
            elif len(summary_x) == 1:
                ## there is no second base
                base2 = "X"
                count2 = 0
            elif len(summary_x) == 2:
                ## there is only one other base
                base2, count2 = summary_x[1]
            elif summary_x[1][1] > summary_x[2][1]:
                ## there is a distinct second top call
                base2, count2 = summary_x[1]
            else:
                ## there is a tie for second top call
                ## get all bases with this count
                tied_x = [tup for tup in summary_x if tup[1] == summary_x[1][1]]
                base2 = "N"
                count2 = sum([c[1] for c in tied_x])
            
            ## save the bases to the chromat variable before possibly changing them to N
            base_chromat = base
            base2_chromat = base2
            ## SIGNAL TO NOISE CHECK
            ## check ratio of top base to second top base
            if count < args.GLOBAL_THRESHOLD_FACTOR * count2:
                ## not enough support.
                base = "N"
            ## implicit else, there is enough support, we use this most frequent base as the call
            
            ## MINIMUM COVERAGE CHECK
            ## check that meets minimum depth threshold
            if count > DEPTH_THRESHOLD:
                consensus += base
                countarr.append(count)
                accuracies.append(100 * (count / sum([c[1] for c in summary_x])))

                ## track second top, for post-hoc inspection.
                consensus_second += base2
                countarr_second.append(count2)

                ## save raw chromat bases
                consensus_chromat += base_chromat
                consensus_second_chromat += base2_chromat
    
    
    ##########################
    ## Step 7: Write output ##
    ##########################
    
    statprint("Writing consensus...")
    with open(args.consensus, "w") as out:
        out.write(">consensus\n")
        out.write("{}\n".format(consensus))

    
    statprint("Writing chromatogram data...")
    with open(args.chromat, "w") as out:
        out.write("pos\tbase\tcount\n")
        for i in range(len(consensus)):
            out.write("{}\t{}\t{}\n".format(i+1, consensus_chromat[i], countarr[i]))
            out.write("{}\t{}\t{}\n".format(i+1, consensus_second_chromat[i], countarr_second[i]))
    
    statprint("Writing per-position consensus accuracies...")
    with open(args.accuracies, "w") as out:
        out.write("pos\taccuracy\n")
        for i in range(len(consensus)):
            out.write("{}\t{}\n".format(i+1, accuracies[i]))

    statprint("Done.")
```
