## Additional File 3 for "Complete sequence verification of plasmid DNA using the Oxford Nanopore Technologies’ MinION device": pseudopair_reads.py

```
TITLE = "PseudoPair Reads"
DESC = "Given a .paf file, find the pseudopaired reads."
'''
@author: sbrown
'''

## Import Libraries

import sys
import argparse
import os
import time


## Declare global variables

DEBUG = False
VERB = False


## Classes and functions

class bcolors:
    CYAN = '\033[1;36;40m'
    BLUE = '\033[1;34;40m'
    GREEN = '\033[1;32;40m'
    YELLOW = '\033[1;33;40m'
    RED = '\033[1;31;40m'
    BOLDWHITE = '\033[1;37;40m'
    DARK = '\033[1;30;40m'
    PURPLE = '\033[1;35;40m'
    ENDC = '\033[0m'

def statprint(msg, msg_type = "STATUS"):
    typeColour = ""
    if msg_type == "ERROR":
        typeColour = bcolors.RED
    elif msg_type == "WARNING":
        typeColour = bcolors.YELLOW
    elif msg_type == "DEBUG":
        typeColour = bcolors.GREEN
    elif msg_type == "SUBPROCESS":
        typeColour = bcolors.GREEN
        msg_type = "     " + msg_type
    else:
        typeColour = bcolors.BOLDWHITE

    print("{message_color}{message_type}{end_color} {time_color}[{datetime}]{end_color}: {message}".format(message_color = typeColour, 
             message_type = msg_type,
             end_color = bcolors.ENDC, 
             time_color = bcolors.BLUE, 
             datetime = time.strftime("%Y/%m/%d %T"), 
             message = msg))


## Main

if __name__ == "__main__":

    ## Deal with command line arguments
    parser = argparse.ArgumentParser(description = TITLE)
    parser.add_argument("--paf", help = "Input paf file", type = str)
    parser.add_argument("--min_align_length", help = "Minimum alignment length to accept", type = int)
    parser.add_argument("--pseudopairs", help = "Output read pairs file", type = str)
    parser.add_argument("-d", "--debug", action = "store_true", dest = "DEBUG", help = "Flag for setting debug/test state.")
    parser.add_argument("-v", "--verbose", action = "store_true", dest = "VERB", help = "Flag for setting verbose output.")
    args = parser.parse_args()

    ## Set Global Vars
    DEBUG = args.DEBUG
    VERB = args.VERB

    print("=======================================================")
    print("Python version: {}".format(sys.version))
    print("Python environment: {}".format(sys.prefix))
    print("Server: {}".format(os.uname()[1]))
    print("Current directory: {}".format(os.getcwd()))
    print("Command: {}".format(" ".join(sys.argv)))
    print("Time: {}".format(time.strftime("%Y/%m/%d %T")))
    print("=======================================================\n")


    ## Script content here

    fwd_reads = {}
    rev_reads = {}

    statprint("Parsing PAF file...")
    for line in open(args.paf, "r"):
        line = line.split("\t")
        read_id = line[0]
        read_len = int(line[1])
        aligned_read_start = int(line[2])
        aligned_read_end = int(line[3])
        strand = line[4]

        aligned_length = aligned_read_end - aligned_read_start

        ## check if read has been seen before. If it has, discard it AND the previous appearance.
        if read_id in fwd_reads:
            del fwd_reads[read_id]
        elif read_id in rev_reads:
            del rev_reads[read_id]
        else:
            ## read not seen before.
            if strand == "+":
                fwd_reads[read_id] = aligned_length
            else:
                rev_reads[read_id] = aligned_length

    statprint("There are {} forward reads and {} reverse reads.".format(len(fwd_reads), len(rev_reads)))

    statprint("Removing reads with short alignments...")
    ## go through and drop read if aligned_length not long enough.
    fwd_to_delete = []
    for ri in fwd_reads:
        if fwd_reads[ri] < args.min_align_length:
            fwd_to_delete.append(ri)
    rev_to_delete = []
    for ri in rev_reads:
        if rev_reads[ri] < args.min_align_length:
            rev_to_delete.append(ri)
    
    for ri in fwd_to_delete:
        del fwd_reads[ri]
    for ri in rev_to_delete:
        del rev_reads[ri]
    
    statprint("There are {} forward reads and {} reverse reads.".format(len(fwd_reads), len(rev_reads)))

    statprint("Pseudopairing...")
    out = open(args.pseudopairs, "w")
    for fr, rr in zip(fwd_reads, rev_reads):
        out.write("{} {}\n".format(fr, rr))
    out.close()

    

    statprint("Done.")
```
